# A Time-calibrated Firefly (Coleoptera: Lampyridae) Phylogeny: Using Genomic Data for Divergence Time Estimation

**DOI:** 10.1101/2021.11.19.469195

**Authors:** Sebastian Höhna, Sarah E. Lower, Pablo Duchen, Ana Catalán

## Abstract

Fireflies (Coleoptera: Lampyridae) consist of over 2,000 described extant species. A well-resolved phylogeny of fireflies is important for the study of their population genetics, bioluminescence, evolution, and conservation. We used a recently published anchored hybrid enrichment dataset (AHE; 436 loci for 88 Lampyridae species and 10 outgroup species) and state-of-the-art statistical methods (the fossilized birth-death-range process implemented in a Bayesian framework) to estimate a time-calibrated phylogeny of Lampyridae. Unfortunately, estimating calibrated phylogenies using AHE and the latest and most robust time-calibration strategies is not possible because of computational constraints. As a solution, we subset the full dataset by applying three different strategies: (i) using the most complete loci, (ii) using the most homogeneous loci, and (iii) using the loci with the highest accuracy to infer the well established *Photinus* clade. The estimated topology using the three data subsets agreed on almost all major clades and only showed minor discordance within less supported nodes. The estimated divergence times overlapped for all nodes that are shared between the topologies. Thus, divergence time estimation is robust as long as the topology inference is robust and any well selected data subset suffices. Additionally, we observed an unexpected amount of gene tree discordance between the 436 AHE loci. Our assessment of model adequacy showed that standard phylogenetic substitution models are not adequate for any of the 436 AHE loci which is likely to bias phylogenetic inferences. We performed a simulation study to explore the impact of (a) incomplete lineage sorting, (b) uniformly distributed and systematic missing data, and (c) systematic bias in the position of highly variable and conserved sites. For our simulated data, we observed less gene tree variation which shows that the empirically observed amount of gene tree discordance for the AHE dataset is unexpected and needs further investigation.

## Introduction

Fireflies (Coleoptera: Lampyridae) consist of more than 2,000 globally distributed described species renowned for their charismatic lighted mating signals. A time-calibrated phylogeny of fireflies would be useful to study their diversification, biogeographical history, and the evolution of their bioluminescence (Fallon et al. 2018; Powell et al. 2021). Furthermore, divergence times on a genus and species level can provide new insights into recent population genetic processes. However, a robust time-calibrated phylogeny of widely-sampled Lampyrids does not currently exist (but see Powell et al. (2021)). The lack of a time-calibrated phylogeny might be surprising given the enigmatic status of fireflies but is possibly due to debated phylogenic relationships (Branham and Wenzel 2001; Stanger-Hall et al. 2007; Martin et al. 2017) and general challenges in dating beetle phylogenies (Toussaint et al. 2017). A recent study by Martin et al. (2019) obtained 436 anchored hybrid enrichment loci (AHEs) for 88 Lampyridae species and 10 outgroup species. In this study, we will use this AHE dataset to estimate a time-calibrated phylogeny of Lampyridae. This study also serves as case-study to evaluate divergence time estimation using genomic data.

Genomic data, such as the AHE dataset by Martin et al. (2019), has promised to solve many outstanding phylogenetic debates (Rokas et al. 2003; Misof et al. 2014; Jarvis et al. 2014). Unfortunately, genomic data has introduced as many or more new challenges. One of the most prevalent problems of phylogenomics is that different genomic datasets (often a method-dependent sub-sample of the genome; Andermann et al. 2020) and inference methods produce conflicting phylogenetic results with high support (Philippe et al. 2017; Betancur-R et al. 2019). Most recent studies have focused on the impact on the inferred tree topology (*e*.*g*., Arcila et al. 2017; Kuang et al. 2018; Alda et al. 2019; Bossert et al. 2021), but other aspects of the phylogenetic inference still need much study. For example, it has been shown that outlier loci (Brown and Thomson 2017; Shen et al. 2017; Walker et al. 2018) and data filtering methods can have a strong impact on the inferred phylogeny. Furthermore, it has been observed that although error in individual gene trees is high *(e*.*g*., Bossert et al. 2021), species tree estimates are often robust. Nevertheless, obtaining robust estimates of gene trees is crucial if we want to study the biological mechanisms underlying gene tree discordance, such as incomplete lineage sorting (Rannala and Yang 2003), gene flow (Hudson et al. 1992) and gene duplication and loss (Rasmussen and Kellis 2012). Finally, much less attention has been paid on estimating divergence time using genomic datasets (but see Smith et al. 2018).

The two most common approaches for inferring (uncalibrated) phylogenies from genomic data are concatenation of all loci and two-step coalescent-based methods. The concatenation method *(e*.*g*., `RAxML` (Stamatakis 2014) and `IQ-TREE` (Minh et al. 2020)) merge all loci together and assume that all loci evolve under the same topology with the same branch lengths. Two-step coalescent-based methods *(e*.*g*., `ASTRAL`, Zhang et al. 2018) estimate first the per-locus gene trees and then estimate the species tree assuming a multispecies coalescent approach. Current two-step approaches are considered superior due to their ability to incorporate incomplete lineage sorting (ILS) but do not provide time-calibrated phylogenies. Therefore, we cannot use a two-step coalescent-based approach to estimate divergence times. The only currently existing methods using sequence data directly to estimate time calibrated phylogenies are full Bayesian coalescent-based methods and concatenation methods for divergence time estimation.

Today, there exists no consensus on estimating a time-calibrated phylogeny using genomic data. Ideally we would like to use all loci and full Bayesian inference methods using relaxed clocks (Drummond et al. 2006). Unfortunately, full Bayesian inference is impossible for genomic datasets due to computational limitations (Harrington et al. 2016; Li et al. 2020). Common approaches include (1) penalized-likelihood methods such as `r8s` (Sanderson 2003) and `treePL` (Smith and O’Meara 2012) (see for example Hamilton et al. 2019; Alda et al. 2019; Opatova et al. 2020; Burbrink et al. 2020); (2) approximate-likelihood methods as implemented in PAML (see for example Harrington et al. 2016; McGowen et al. 2020; Li et al. 2020), and (3) full-likelihood Bayesian divergence time analysis using a relaxed clock model, as implemented in `BEAST` (Drummond et al. 2012) and `RevBayes` (Höhna et al. 2016), on a subset of the available data (see for example Harrington et al. 2016; Ericson et al. 2020; Bianconi et al. 2020).

Penalized likelihood approaches are faster to compute but do not use the sequence data directly. Thus, penalized likelihood approaches are less robust because they do not fully take the uncertainty in branch length estimates into account (Ho and Duchêne 2014). Approximate-likelihood methods are also faster than full-likelihood methods but their accuracy has not been compared against another. Since full-likelihood Bayes divergence times methods are most widely used and well established, we focus on and explore the third option. Specifically, we will focus on different approaches to sub-sample the full AHE dataset. Several approaches to subsample the full dataset have been proposed: (1) choose a random subset of the loci (Harrington et al. 2016; Alda et al. 2019; Ericson et al. 2020), (2) choose the most complete loci (Harrington et al. 2016), and (3) choose the loci with lowest GC variation (Romiguier et al. 2013). Additionally to the second and third option, we selected loci with a high phylogenetic accuracy to recover the established genus *Photinus*. Thus, before estimating the divergence time using the three concatenated data subsets, we explored each single AHE locus to identify reliable loci and exclude outlier loci (Brown and Thomson 2017; Walker et al. 2018). We estimated the phylogeny using each AHE locus individually, producing 436 posterior distributions of phylogenetic trees. We used these individual phylogenies to (1) explore the gene-tree discordance, (2) support for named sub-families, tribes and genera, (3) correlation between data summary statistics and gene tree error. Finally, we performed a simulation study to test the impact of (a) incomplete lineage sorting, (b) uniformly distributed and systematic missing data, and (c) systematic bias in the position of highly variable and conserved sites. All of the methods described in this paper have been implemented in the Bayesian phylogenetic inference software package `RevBayes` (Höhna et al. 2016).

## Methods and Data

### Lampyridae Anchored Hybrid Enrichment (AHE) Dataset

In this study we used the 436 anchored hybrid enrichment sequences from Martin et al. (2019). The dataset contains 88 Lampyridae species and 10 outgroup species. The AHEs have been trimmed and cleaned (Martin et al. 2019). Martin et al. (2019) kept only loci with an overall sequence completeness of 50%. Here we use exactly the same alignments downloaded from doi:10.5061/dryad.737c8t8.

Our specific focus in this study was the genus Photinus. Photinus is the second most specious genus of Lampyridae comprising of ∼ 240 species with a Neartic and Neotropical distribution (McDermott 1964). Only in the past 15 years, more than 40 new Photinus species have been described (Zaragoza-Caballero 2007, 2015; Zaragoza-Caballero et al. 2020). It is hypothesized that the origin of Photinus resides in Tropical America (McDermott 1964) and the estimation of the age of this genus will be the first step into elucidating its biogeographical history.

Fireflies from the genus *Ellychnia* were traditionally placed outside *Photinus* but recent molecular studies have shown that *Photinus* is paraphyletic and *Ellychnia* is grouped within it (Stanger-Hall et al. 2007; Lower et al. 2017; Martin et al. 2017). From a morphological perspective, *Ellychnia* and *Photinus* share morphological characteristics that support placing these into the same genus (Zaragoza-Caballero 2012, 2015; Zaragoza-Caballero et al. 2020). Thus, we consider the combined clade of *Photinus* and *Ellychnia* in this study.

#### Sequence coverage

First, we explored the sequence coverage of the 436 sequence alignments. Figure S1 shows the percentage of missing sites per taxon and AHE locus. That is, we specifically looked whether a given taxon (*i*.*e*., column) or locus (*i*.*e*., row) had considerable lower sequence coverage (white or light gray). The distribution of missing sites is not homogeneous and particular taxa are more affected than others. Sequences for *Photinus* and *Ellychnia* species had a comparably high sequence coverage.

#### Data summary statistics

We computed several summary statistics for the data which might indicate the usefulness of each AHE locus. The summary statistics were: (1) number of variable sites, (2) number of invariable sites, (3) minimum pairwise distance between any taxon pair, (4) maximum pairwise distance between any taxon pair, (5) minimum GC content of any taxon for this locus, (6) maximum GC content of any taxon for this locus, (7) average GC content for this locus, (8) variance in GC content over all taxa for this locus.

The number of variable sites should be a predictor for the informativeness of the locus, with the expectation that loci with a higher number of variable sites have more information to resolve the phylogeny correctly. Conversely, the number of invariant sites should be lower to obtain more phylogenetic information. Nevertheless, it might be the case that if all sites are variable (i.e., no invariant sites), then the locus is likely saturated and most information is lost due to multiple substitutions. Alternatively, we could use the fraction of variable sites to invariant sites, although this fraction is only informative in the context of the sequence length. Therefore we used the number of variable and invariant sites directly to avoid redundancy.

The minimum pairwise distance shows how well we can expect to resolve the phylogeny on a species level. If there are some species with identical sequences for this AHE locus, then we have no information about the divergence between the species except that they should be placed very closely together. The maximum pairwise distance can indicate if there are outlier sequences which bias our phylogenetic reconstruction. If such outlier sequences with high pairwise distance to all other sequences exist, then this indicates non-orthologous sequences or miss-alignments which will lead to wrong placements of the taxa in the inferred phylogeny.

The GC content has been suggested to be an indicator of gene tree error with GC rich loci having a higher error (Romiguier et al. 2013). A high GC content is hypothesized to be correlated with high rates of recombination through GC-biased gene conversion (Mugal et al. 2015) and therefore these regions can be problematic for phylogenetic inference (Romiguier et al. 2013). Similarly, a high variance in GC content could indicate branch heterogeneous or non-stationary substitution processes (Betancur-R et al. 2013), for example due to convergent evolution which would also bias phylogenetic inference.

### Exploration of Individual AHE Loci

#### Inference of phylogenies per AHE locus

We performed a phylogenetic analysis for each individual locus. The goal was to estimate the tree topology and species relationships without confounding factors of molecular clocks and divergence times. Therefore, we performed a standard Bayesian phylogenetic analysis, which had been shown recently to perform best for AHE loci (Bossert et al. 2021). First, for single loci we removed sequences with less than 50% sequence coverage to avoid problems in placing poorly sequenced and possibly degraded samples. Our analysis model consisted of a GTR+Γ+I substitution model (Tavar’e 1986; Yang 1996) and a uniform prior distribution on tree topologies with an exponential prior distribution on the branch lengths (Höhna et al. 2017). We ran four replicated MCMC runs with 50,000 iterations each (with, on average, 167.8 moves per iteration). We sampled phylogenies every iteration.

#### Posterior support of known clades

For each AHE locus, we computed the posterior probability of 26 named subfamilies, tribes and genera (see Table S1). We only computed the posterior probability for a clade if the locus contained at least two species for that clade. Specifically, we computed the posterior probability if the given clade was found to be strictly monophyletic according to known classifications. The posterior probabilities show us (a) which known clades are supported, and (b) how much variation in support exists for the known clades between loci. We expect that well established clades, such as *Photinus + Ellychnia*, should overall be well supported. Nevertheless, we would not be surprised to see some variation in support as gene trees are expected to be different from species trees (Maddison 1997). For example, the multispecies coalescent process predicts that gene trees can be different to the species tree if internal branches are very short and population sizes are very large (Rosenberg and Tao 2008; Huang and Knowles 2009). However, the discordance between species tree and gene trees should be restricted to local difference within few coalescent units (Degnan and Rosenberg 2009) and not produce gene trees that are drastically different from the species tree.

#### Model Adequacy Testing

Additionally, for each locus we performed posterior prediction simulations to check for model adequacy using the *P* ^3^ pipeline (Höhna et al. 2018). Posterior predictive distributions are used to perform model adequacy testing, *i*.*e*., testing the *absolute* fit of a model to the observed data (Bollback 2002). If the model shows a bad absolute fit to the data, then estimates, such as the tree topology, can be biased (Brown 2014). For example, if our model predicts much lower variation in GC content among sequences, then our inference might wrongly group taxa with low (or high) GC content together (Betancur-R et al. 2013).

Posterior predictive distributions were simulated using parameters values (*e*.*g*., phylogeny and substitution rates) drawn from the posterior distribution which were inferred in the previous step. We conservatively discarded the initial 50% of samples as burnin and used the remaining 100,000 samples for the simulations (four replicates with originally 50,000 samples each). Finally, we computed the posterior predictive p-values as the frequency how often the summary statistic of the observed data was larger or equal to the summary statistic computed using the simulated data (midpoint p-values; Höhna et al. (2018)). That is, if we obtain a very low p-value, then most or all of our simulated datasets have a larger summary statistic. For example, if our empirical alignment had very few variable sites and most simulated datasets had more variable sites, then the p-value would be close to zero. Conversely, a high posterior predictive p-value depicts larger summary statistics from the observed data compared with the simulated data.

### Simulation Study

We performed a simulation study as a benchmark and reference for our single locus phylogenetic analyses. Specifically, we focused on (1) the discordance between gene trees and species trees under the multispecies coalescent model, (2) the impact of missing sequence data on phylogeny inference and model adequacy testing, and (3) the impact of unequal distribution of fast versus slow evolving sites in combination with missing sequence data on phylogeny inference and model adequacy testing. First, under the multispecies coalescent model we expect that gene trees differ from the species to some extend purely due to the stochastic process (Degnan and Rosenberg 2009). For example, assuming a population size of 100,000 diploid individuals and a generation time of one year, the expected time of a coalescent event between two individuals is 200,000 years. Then, if the branch leading to the next speciation event is shorter than the coalescent time between two individuals, then we could observe deep coalescent events with incomplete lineage sorting. Thus, to observe incomplete lineage sorting the population size needs to be sufficiently large and/or the internal branch length needs to be sufficiently short. In our simulations, we simulated 436 gene trees within the fixed species tree (see below) and three different population sizes: 100,000 diploid individuals, 1,000,000 diploid individuals and 10,000,000 diploid individuals. The chosen population sizes for the simulations were based on known insect effective population sizes (Keightley et al. 2015; Crossley et al. 2019; Arguello et al. 2019; Kapopoulou et al. 2020) and were conservatively large to explore the potential effect.

Second, the AHE dataset —as most phylogenomic datasets— are far from complete and missing sequence data is heterogeneously distributed (see Figure S1). On the one hand, missing sequence data can impact phylogeny inference, specifically if some taxa have a high fraction of missing sequence data (Sanderson et al. 1998; Lemmon et al. 2009). These taxa are often rogue and cannot be placed with certainty or correctly in the phylogeny (Thomson and Shaffer 2010). On the other hand, several simulation studies have shown that if missing data are homogeneously distributed or the number of informative sites is large, then missing data are not problematic (Wiens 2003; Roure et al. 2013). Much less attention has been given to model adequacy testing and computing summary statistics with missing sequence data. For example, if a given site (*i*.*e*., column) in the alignment contains mostly missing sites but the few actual sites are identical, it is then unclear if this site is invariant or not. Thus, missing sites can impact our calculation of summary statistics, and thus our evaluation of model adequacy. Here, we explore the impact of missing data with a specific focus on how missing data is distributed in AHE datasets.

We simulated sequence alignments for each of the three sets of 436 gene trees as follows. We simulated branch rates from a uncorrelated lognormal relaxed clock model with mean 1.836 × 10^−3^ (in million years) and standard deviation of 0.58. Then, we simulated sequence data under a GTR+Γ model with base frequencies π = {0.31, 0.17, 0.19, 0.33}, substitution rates ϵ = {0.087, 0.295, 0.08, 0.09, 0.38, 0.068} and site rate categories *r* = {0.039, 0.271, 0.841, 2.849}. The lengths of the sequences was determined from the corresponding empirical alignment. All values were retrieved from the empirical concatenated analysis to provide biologically realistic simulation settings. Additionally, each simulated alignment was masked so that the same positions in the data matrix were missing for both the empirical dataset and simulated dataset. This procedure to create alignments with missing data by applying masks obtained from the empirical alignments produces patterns where missing data are not uniformly distributed but clustered around the beginning and end of the alignment as well as on given taxa (see Supplementary Figures S2-S4). Thus, we obtained two sets of alignments for each simulated alignment (with and without missing sites).

Third, the AHE dataset has a heterogeneous distribution of variable sites where most variable sites are at the flanking regions and most invariant sites are at the center of the locus (Faircloth et al. 2012). This heterogeneous distribution of fast versus slow evolving sites stands in strong contrast to the model assumption of standard phylogenetic models. The among site rate variation model (+Γ) allows for rate variation using four discrete rate categories but each site evolves independently and identically distributed. That is, each site has a probability of 0.25 to be in any of the four rate categories regardless of the position in the alignment (center vs. beginning/end). This model violation might not be a problem for many phylogenetic analyses. However, the combination of missing data that is more prevalent at the same positions as highly variable sites could induce a systematic bias. We explored this potential systematic bias by repeating the above simulation with rate categories drawn deterministically depending on the position in the alignment. Specifically, we divided the alignment in eight equal-sized regions where the outer regions received the highest of the four rate categories and the middle regions the lowest rate categories respectively.

In total, we simulated three sets of 436 gene trees and four alignments per gene trees (436 loci x 3 population sizes x 2 levels of missing data x 2 modes of rate variation = 5232 simulated alignments). We analyzed each simulated alignment with the same inference pipeline as the empirical AHE dataset. We performed an MCMC analysis for each alignment, a posterior predictive simulation, and computed the posterior predictive p-values and posterior probabilities of the pre-defined clades.

### Divergence time estimation of Lampyridae phylogeny

We used the 436 ultra-conserved elements (AHEs) recently published by Martin et al. (2019) for the Lampyridae divergence time estimation. Because of computational limitations we could not perform a phylogenetic analysis on all 436 AHE loci jointly with a model of appropriate complexity (*e*.*g*., each AHE loci having its own unlinked GTR+Γ substitution model). Instead, we selected three data subsets (Figure 1). The first data subset contained all loci with at 95% average sequence coverage (Figure 1) because gappy sequences (*i*.*e*., low sequence coverage) could indicate sequencing and/or alignment problems. Additionally, missing data reduce information in the alignment (Philippe et al. 2004) and we aimed to maximize the phylogenetic information for the associated computational cost. The second data subset contained all AHE loci which supported the genus *Photinus + Ellychnia* to be monophyletic because we constrained *Photinus + Ellychnia* to be monophyletic for the fossil calibration (Table 1). Using loci that conflict with the enforced calibration constraint could lead to biased results (Yang and Rannala 2006). The third data subset consisted of all AHE loci with low variation in GC content among taxa. Increased variance in GC content among taxa is often a signal of compositional heterogeneity which is not modeled appropriately using standard phylogenetic substitution models (*e*.*g*., GTR+Γ) and can lead to wrong phylogenetic inferences (Foster 2004; Romiguier et al. 2013; Betancur-R et al. 2013; Duch^ene et al. 2017). The data subsets contained 37, 24 and 30 AHE loci for the high sequence coverage, *Photinus + Ellychnia* monophyly, and low GC variance criteria respectively. These data subsets share some loci but the majority of loci are private for each data subset (Figure 1). Thus, our divergence time analyses using these three different data subsets are mainly independent. If all three datasets produce the same or highly similar divergence time estimates, then we are confident that the divergence time estimates are robust to our choice of data subsets.

**Table 1:**
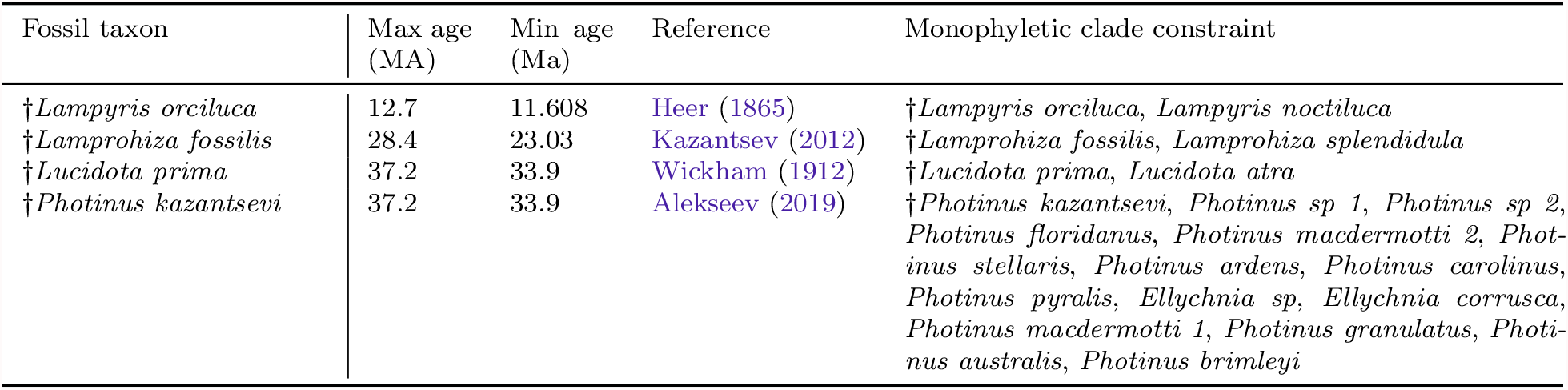
Fossils and calibration constraints for the time-calibrated divergence time analysis.

**Figure 1:**
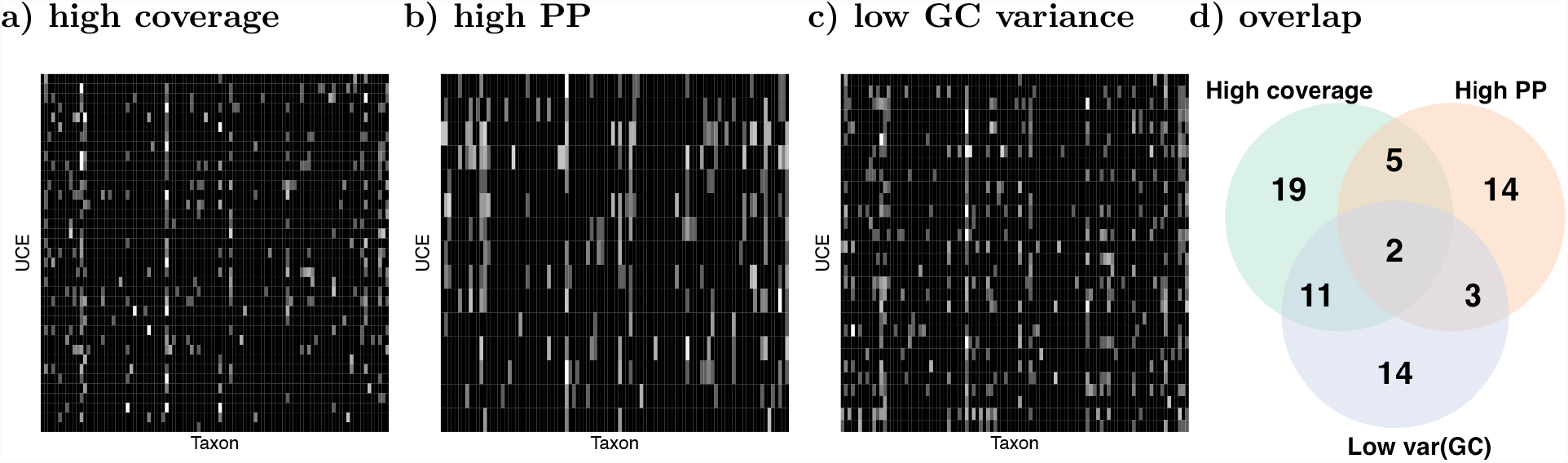
Overlap and completeness of the AHE data subsets. We selected three data subset based on (a) loci with on average 95% completeness (high coverage), (b) loci with a high posterior probability of *Photinus + Ellychnia* being monophyletic (high PP), and (c) loci with low variance in GC content. (a–c) shows the completeness of the selected loci. We computed the percentage of sites missing per sequence. Black cells depict complete sequences and white cells depict entirely missing sequence. The gray shades depict the percentage in between. Each row represents one of the AHE loci and each column represent one taxon. (d) shows the overlap between data subsets. Despite there being some overlap between the data subsets, the majority of loci is private to each subset.

For each data subset we employed a partitioned GTR+Γ substitution model (Tavar’e 1986) where among site rate variation was modeled by 4 discrete categories obtained from a gamma distribution (Yang 1994). We did not perform any substitution model selection (*e*.*g*., Tagliacollo and Lanfear 2018*) as Bayesian inference is robust to substitution model over-parametrization (Huelsenbeck and Rannala 2004; Lemmon and Moriarty 2004; Abadi et al. 2019). Thus, our chosen substitution model is conservative albeit computationally more demanding because it assigns each partitions its own set of substitution model parameters. We applied standard prior distributions for the substitution model parameters, that is, a flat Dirichlet prior distributions on both the stationary frequencies and on the exchangeability rates (Höhna et al. 2017). Furthermore, to account for rate variation among lineages we used a relaxed-clock model with uncorrelated lognormal distributed rates (UCLN, Drummond et al. 2006). We applied an uninformative hyperprior distribution on both the mean ∼ uniform(0, 100) and standard deviation, sd ∼ uniform(0,100) of the branch-specific clock rates*.

*Estimating a dated phylogeny of most insect clades is extremely challenging because of the lack of appropriate fossils for node calibrations. We used four fossil taxa belonging to different genera within Lampyridae (Table 1). We used †Lampyris orciluca* (Heer 1865) which belongs to the genus *Lampyris*, and †*Lucidota prima* (Wickham 1912) which belongs to the genus *Lucidota*. Additionally we used †*Lamprohiza fossilis* (Kazantsev 2012) and placed it belonging to the genus *Lamprohiza*, although †*Lamprohiza fossilis* was previously named †*Phausis fossilis* (Beier 1952). Since *Phausis* and *Lamprohiza* are the only two genera in the subfamily Lamprohizinae, our calibration is equivalent to a minimum age of Lamprohizinae because the split *Phausis* – *Lamprohiza* must have occurred before the age of †*Lamprohiza fossilis*.

We included the recently published fossil for the *Photinus* clade; †*Photinus kazantsevi* found in Baltic amber and dated to the Upper or Mid-Eocene (33.9 to 47.8 ma, Alekseev 2019). However, the taxonomic placement of this fossil specimen is unknown, *i*.*e*., whether this fossil represent a stem or crown fossil. To explore the sensitivity of our fossil calibrations and divergence time analyses, we performed each divergence time analyses for the three data subsets twice; once including †*Photinus kazantsevi* and enforcing *Photinus* to be monophyletic and the other time excluding †*Photinus kazantsevi*. This sensitivity analysis provides both insights into the robustness of the divergence time analysis when a fossil is excluded (Near and Sanderson 2004; Saladin et al. 2017) and the placement of †*Photinus kazantsevi*.

We omitted using the fossil †*Electrotreta rasnitsyni* because our preliminary analyses showed that *Drilaster sp* and *Stenocladius shirakii* were not recovered as sister species (see also Martin et al. 2019). We did not want to enforce the sister relationship between *Drilaster sp* and *Stenocladius shirakii* because this could bias the phylgeny inference. Similarly, we omitted the use of the fossil †*Protoluciola albertalleni* (Kazantsev 2015) because the phylogenetic relationship of *Luciola* are under continuous revision and thus the placement of the fossil is very uncertain. Nevertheless, since †*Protoluciola albertalleni* is a fossil Luciolinae from Burmite amber (∼99 million year old), it provides a mimum age for Lampyridae of at least 99Ma.

We used the fossilized birth-death-range process (Stadler et al. 2018) to time-calibrate the Lampyridae phylogeny. The fossilized birth-death range process requires assignment of fossils to clades (see Table 1) and integrates over both the actual placement within the clade (*e*.*g*., stem vs crown) and the actual time of fossil within the specified stratigraphic range. That is, we provided both minimum and maximum ages (Table 1) for each fossil taxon. Then, the fossilized birth-death range process gives equal probability that the true age of the fossil was within the specified range. In principle, we could omit the monophyletic constraints if we had morphological data for both fossil and extant taxa using tip-dating approaches (Ronquist et al. 2012; Arcila et al. 2015; Gavryushkina et al. 2017). Unfortunately, there does not exist an appropriate morphological dataset for fossil and extant Lampyridae which currently prohibits tip-dating approaches.

Estimating the divergence times under a relaxed-clock model is extremely challenging because of the non-identifiability between evolutionary rates and time (Donoghue and Yang 2016). We used a newly developed MCMC move, the `RateAgeBetaShift`, to alleviate the problem of highly correlated parameter estimates (Zhang and Drummond 2020). Additionally, we performed 12 independent Metropolis-Coupled MCMC (MCMCMC, Altekar et al. 2004) runs with one cold and seven heated chains for 50,000 iterations (with on average 458 moves per iteration) using the parallelized version of `RevBayes` (Höhna et al. 2021). Each single MCMCMC replicate took up ∼1, 111, ∼618 and ∼956 hours (for the three data subsets respectively) using 8 CPUs simultaneously with a total of ∼515, 710 CPU hours (∼21, 487.93 CPU days or ∼58.87 CPU years). This high computational cost using only 24 to 37 loci demonstrates that it is computationally unfeasible to perform joint Bayesian divergence time analyses using all 436 loci.

## Results

### Properties of the AHE loci

We obtained a minimum of 218 variables sites and a maximum of 2,071 variable sites with a mean of 696 variable sites (Figure S5). Similarly, we obtained a minimum of 23 invariable sites, a maximum of of 1067 invariable sites and a mean of 225 invariable sites. We used the number of variable sites as a proxy for how informative a locus is (Townsend 2007). Overall, the distribution of the number of variable sites appeared unimodal without extremely low outliers. Thus, we did not see any indication that specific loci should be particularly poor for phylogenetic inference.

The minimum and maximum pairwise distance showed interesting patterns. The majority of loci had a minimum pairwise distance of zero (Figure S5), which means that the alignments contained two sequences without substitutions among them. Hence, there is no phylogenetic signal to distinguish between the sequences. In itself, this low pairwise distance does not imply a problem for phylogenetic inference because the two taxa will be placed as sister taxa. However, this distribution could indicate that there are several taxa that cannot be resolved.

The maximum pairwise distance showed a skewed distribution with some larger outliers. This could indeed be problematic. First, the high maximum pairwise distance will most likely lead to long branches in the phylogeny. Second, the high distance could occur due to non-homologous sequences. The sequences, for example, could be contaminated, mis-aligned and/or represent paralogs.

The distribution of GC content showed some slightly multi-modal and skewed mean GC content and variance in GC content (Figure S5). The mode with lower mean GC content and higher variance in GC content could represent loci which are problematic for phylogenetic inference.

### Gene trees

#### Posterior probabilities of named clades

Our single loci (gene trees) phylogenetic analyses yielded very mixed results (Figure 2 and S6). On a subfamily level, monophyly of Lampyrinae and Amydetinae was rejected by all 436 loci, whereas the monophyly of Luciolinae and Photurinae was rejected by the majority of loci (Figure S6). The monophyly of Ototretinae was ambiguously supported and Lamprohizinae was the only subfamily which we recovered as monophyletic, which could be due to taxon sampling (i.e., Lamprohizinae was represented by only two taxa). The results on a tribe level were similar; either all or the majority of loci rejected the monophyly of all six tribes (Cratomorphini, Lamprocerini, Lampyrini, Photinini, Phosphaenini and Luciolini; Figure S6). The monophyletic support increased on the genus level; eight of the fourteen genera were recovered as monophyletic, five genera were rejected as being monophyletic and *Australoluciola* was ambiguously recovered as either monophyletic or not (Figure S6).

**Figure 2:**
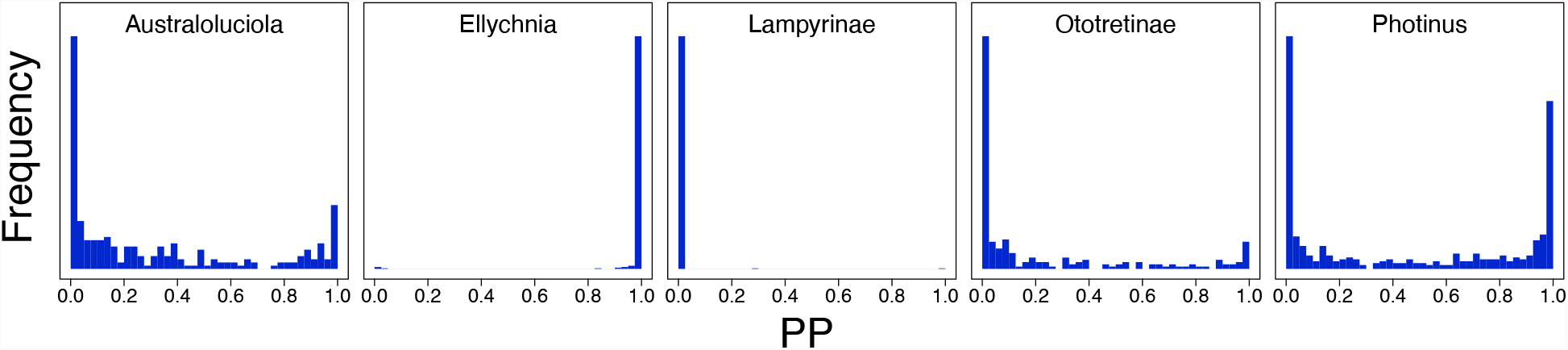
Posterior probability of being monophyletic per AHE locus for five example clades (3 genera and 2 subfamilies). For each AHE locus, we computed the posterior probability that the clade is monophyletic. A histogram with most loci having a high posterior probability (*e*.*g*., *Ellychnia*) depicts strong support by the majority of AHE loci. Conversely, a histogram with most loci having a low posterior probability (*e*.*g*., *Ototretinae*) depicts strong support against monophyly by the majority of AHE loci. Other clades (*e*.*g*., *Photinus*) received contradicting support with some loci strongly supporting and other loci strongly rejecting monophyly. Note that we assumed *Photinus* being paraphyletic and instead considered *Photinus+Ellychnia* based on previous results. The histograms for all clades is shown in the supplementary material Figure S6.

#### Correlation between missing data and posterior support

Given the poor support on higher taxonomic levels, and the ambiguous support for some of the named clades, we investigated whether there is a correlation between missing data and phylogenetic accuracy. Here, we associate phylogenetic accuracy with the ability to recover monophyly of an established clade. Specifically, we used the posterior probability of *Photinus+Ellychnia* being monophyletic. We focus here on *Photinus+Ellychnia* because it is a well studied genus whose monophyly is not debated (Stanger-Hall et al. 2007; Lower et al. 2017; Martin et al. 2019) although we observed rather ambiguous support (Figure 2). The same investigation for all named clades is shown in the Supplementary Material Figure S7.

We observed that there is no correlation between sequence coverage and the posterior probability of *Photinus+Ellychnia* being monophyletic (Figures S7 and S8). This results is actually expected because we removed taxa which had 50% or more sites in the sequence missing. The overall sequence coverage instead represents the completeness of the entire alignment and therefore loci with higher average sequence coverage are loci that contain more taxa after pruning. The same trend and correlation between sequence coverage and phylogenetic accuracy can be seen for all other tested clades (Figure S7). Thus, our pruning of incomplete sequences from the alignment makes filtering loci based on overall sequence coverage futile.

#### Correlation between summary statistics of the data and posterior support

Next to the sequence completeness of a loci, other summary statistics of the data could provide good indicators about the quality and usefulness of a locus (phylogenetic accuracy). Again, we used the ability to recover monophyly of the clade *Photinus+Ellychnia* as a predictor for phylogenetic accuracy. We compared several summary statistics to the posterior probability of *Photinus+Ellychnia* being monophyletic (Figure 3, green dots and green dashed line, and Figure S8). Instead of seeing clear trends (*i*.*e*., monotonously increasing or decreasing correlations), we observed unimodal correlation (*e*.*g*., for the number of invariant sites). That means that outlier loci with extreme values for the summary statistics produce lower phylogenetic accuracy (*e*.*g*., number of invariant sites, minimum GC content and maximum GC content). Only for the minimum pairwise distance and the variance in GC content did we observe a positive correlation (and negative correlation, respectively) with phylogenetic accuracy. A negative correlation between the variance in GC content and phylogenetic accuracy is expected because a low variance in GC content corresponds to more homogeneous substitution processes which are easier to model and produce less biased phylogenetic estimates (Foster 2004). Most importantly, the single loci (Figure 3, green dots) show large variations and the overall trends computed by the average might mislead conclusions.

**Figure 3:**
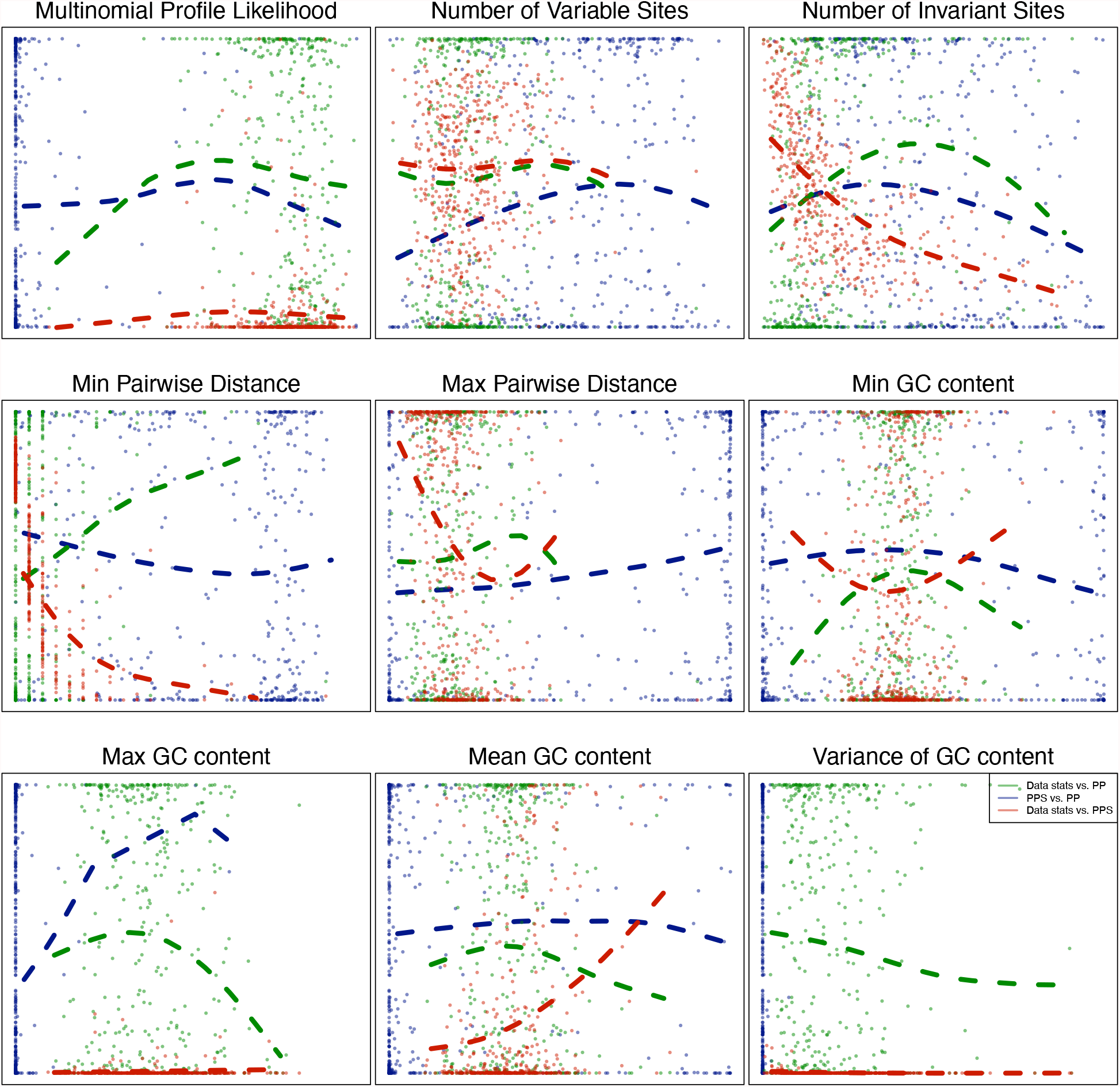
Comparison between phylogenetic accuracy (*i*.*e*., posterior probability of the clade *Photinus+Ellychnia* being monophyletic), model adequacy (*i*.*e*., posterior predictive p-values) and data summary statistics obtained for the AHE dataset of Martin et al. (2019). In green we show the comparison between data summary statistics (x-axis) and the posterior probability of *Photinus* being monophyletic (y-axis). In blue we show the comparison between posterior predictive p-values (x-axis) and the posterior probability of *Photinus* being monophyletic (y-axis). In red we show the comparison between data summary statistics (x-axis) and posterior predictive p-values (y-axis). The dashed lines represent smoothed spline function of the corresponding comparisons. Note that for the variance of GC content we could not plot a predictor function because all posterior predictive p-values are 0.0. We observe that there is no correlation between model adequacy (posterior predictive p-values) and the posterior probability of *Photinus+Ellychnia* being monophyletic (blue line). Interestingly, we observe some correlation between data summary statistics and model adequacy (red line). For each AHE locus, we computed the posterior probability that the clade *Photinus + Ellychnia* is monophyletic. In the supplementary material we show each comparison separately (Figure S8, S10 and S11).

#### Model adequacy

Our posterior predictive simulations showed clear model violations for all loci (Figure S9). No single locus passed all eight posterior predictive checks using a significance level of α = 0.05. Thus, based on our pipeline and our model adequacy checks, we do not have an appropriate phylogenetic model for a single locus. All of our gene tree estimates could be biased due to model violations. If we would filter our original dataset based on which locus passed all model adequacy checks, then we would be left without any locus to proceed further.

The posterior predictive results for the minimum, maximum, mean and variance of GC content are difficult to interpret. The minimum GC content of the posterior predictive datasets was either too low (posterior predictive p-value close to 0.0) or too high (posterior predictive p-value close to 1.0) for the majority of loci. Since our phylogenetic substitution model assumes a homogeneous process with all sequences having the same stationary distribution (*i*.*e*., same expected GC content), it is possible that we do not correctly model outliers sequences with either high or low GC content. This hypothesis corroborated that our posterior predictive datasets have too low variance in GC content (Figure 5, posterior predictive p-value close to 0.0). Nevertheless, it is unexpected that our posterior predictive datasets have too low mean GC content. The mean GC content should be modeled accurately by the stationary distribution of the substitution process.

We observed no clear correlation between the posterior predictive p-values and the gene tree estimation accuracy (when assuming monophyly of *Photinus+Ellychnia* as a proxy for gene tree accuracy, Figure 3 blue dots and dashed blue line). However, we observed a negative correlation between several summary statistics and posterior predictive p-values (Figure 3 red dots and dashed red line). This indicates that large summary statistics represent loci which are likely to be outliers which we cannot model adequately. For example, a high minimum or maximum pairwise distance could be alignment errors and removing these loci could improve phylogenetic inference.

### Simulation Study

In our simulation study, we simulated 12 sets of 436 loci under different conditions. The motivation of the simulation study was to establish (1) how much gene tree error is realistic, (2) the impact of missing sequence data on phylogeny inference and model adequacy testing, and (3) the impact of unequal distribution of fast versus slow evolving sites in combination with missing sequence data on phylogeny inference and model adequacy testing.

First, we observed that simulated complete alignments had different distributions of summary statistics compared to the empirical data (Figure 4). Interestingly, when we masked the alignments to mimic the distribution of missing sequence data as in the original AHE dataset, then we obtained comparable summary statistics. Specifically, the distribution of minimum and maximum pairwise distance matched the empirical distribution only if we removed sites distributed exactly as in the empirical dataset. Similarly, the distribution of mean and variance of GC content matched between simulated dataset and empirical dataset only if we removed sites distributed exactly as in the empirical dataset. Thus, the observed variance in GC content from the empirical data could be a bias observed due to the missing data. However, the distribution of maximum and minimum GC content are wider for the empirical data than for the simulated data. Therefore, not all aspects of the empirical data could be explained solely due to missing data. Furthermore, the systematic distribution of highly variable sites at the beginning and end of the sequences compared with the conserved regions in the center did not impact the computed summary statistics.

**Figure 4:**
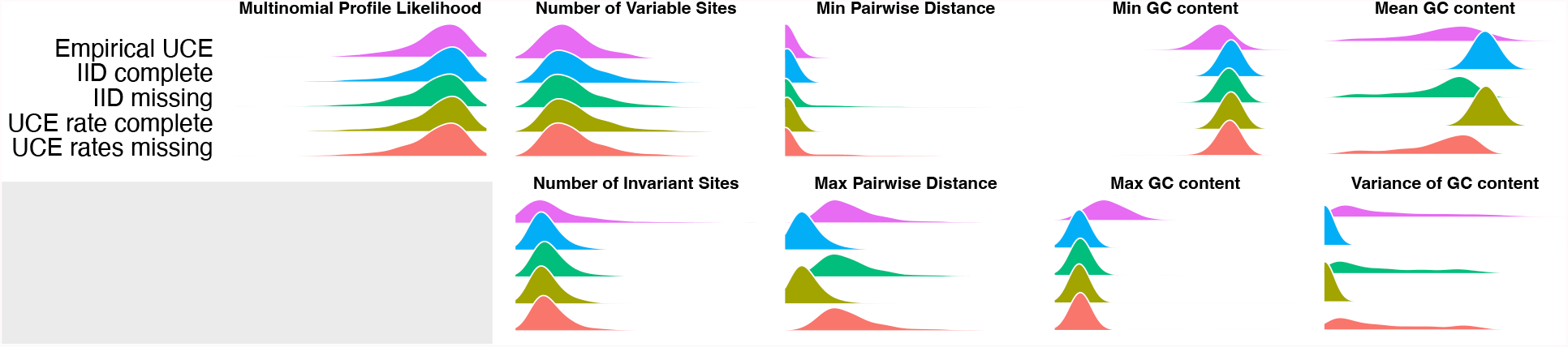
Summary statistics of the empirical AHE datasets and simulated datasets. The bottom row shows the summary statistics computed for the empirical dataset of Martin et al. (2019). The simulated datasets show similar distributions to the empirical dataset only if missing sequences were considered. The second row shows the summary statistics computed for the simulated dataset with complete sequences and homogeneous (*i*.*e*., independent and identically distributed, IID) highly variables sites. The third row shows the summary statistics computed for the simulated dataset with missing sequences and homogeneous highly variable sites. The fourth row shows the summary statistics computed for the simulated dataset with complete sequences and systematically distributed (*i*.*e*., akin to the empirical AHE dataset) highly variables sites. The bottom row shows the summary statistics computed for the simulated dataset with missing sequences and systematically distributed highly variable sites. The distribution of highly variables versus conserved sites had little to no impact on the summary statistics.

**Figure 5:**
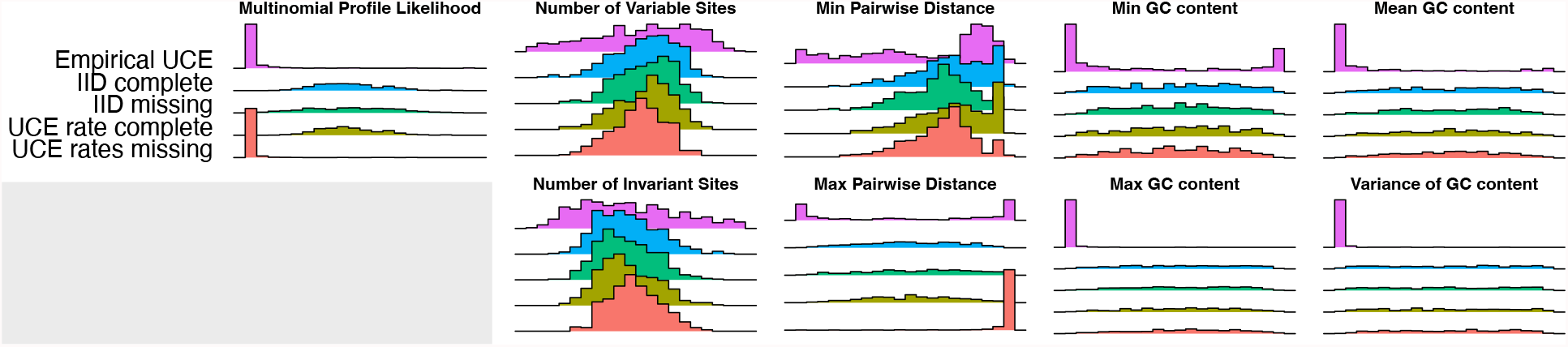
Posterior predictive p-values for the empirical and simulated datasets. The top row shows the frequency of posterior predictive p-values for the empirical dataset of Martin et al. (2019). The second row shows the posterior predictive p-values for the simulated dataset with complete sequences and homogeneous (*i*.*e*., independent and identically distributed, IID) highly variables sites. The third row shows the posterior predictive p-values for the simulated dataset with missing sequences and homogeneous highly variable sites. The fourth row shows the posterior predictive p-values for the simulated dataset with complete sequences and systematically distributed (*i*.*e*., akin to the empirical AHE dataset) highly variables sites. The fifth row shows the posterior predictive p-values for the simulated dataset with missing sequences and systematically distributed highly variable sites. The empirical dataset has mostly posterior predictive p-values of either 0.0 or 1.0, indicating model violation and inadequacy. Conversely, non of the simulated datasets showed model violations as the model used for simulation and inference was identical. Missing data did not impact the posterior predictive p-values and thus the computation of the summary statistics in a systematically biased way.

Our posterior predictive simulations using the simulated data showed that our phylogenetic model was adequate, except in the case when the data were simulated with highly variable sites at the ends of the alignment and sites missing from the alignments. The observed good model adequacy is not surprising because the model used for simulation and inference matched but instead very reassuring that our implementation of the models is indeed correct. Furthermore, our results imply that missing data, when distributed in a systematic manner, can bias our posterior predictive p-values and model adequacy tests. Specifically, we found that when using the the multinomial likelihood and the maximum pairwise distance for the simulated data with the combination of systematically ordered highly variables sites at the borders and missing sequence data, then our model adequacy tests for the simulated failed. It remains surprising that our phylogenetic substitution model was not adequate for even a single empirical locus.

We observed an unexpected amount of gene tree discordance between our species tree and the gene trees (Figure 6). The RF-distance computed for the empirical dataset are more centered at intermediate values and never close to 0. That means, not a single gene tree was equal to or close to any of our reference trees. Using the simulated data as a reference predicts that we should observe more often gene trees that are similar to the species tree. Only if we assumed an unrealistic large population size of 10,000,000 diploid individuals did we observe gene tree discordance for the simulated data close to the empirical data (Figure S12). For such large population sizes, the coalescent units are 20 million years, which means that deep coalescent events are probable to occur even 100 million years before the species diverge. It remains elusive to what reason is causing this unexpected gene tree discordance.

**Figure 6:**
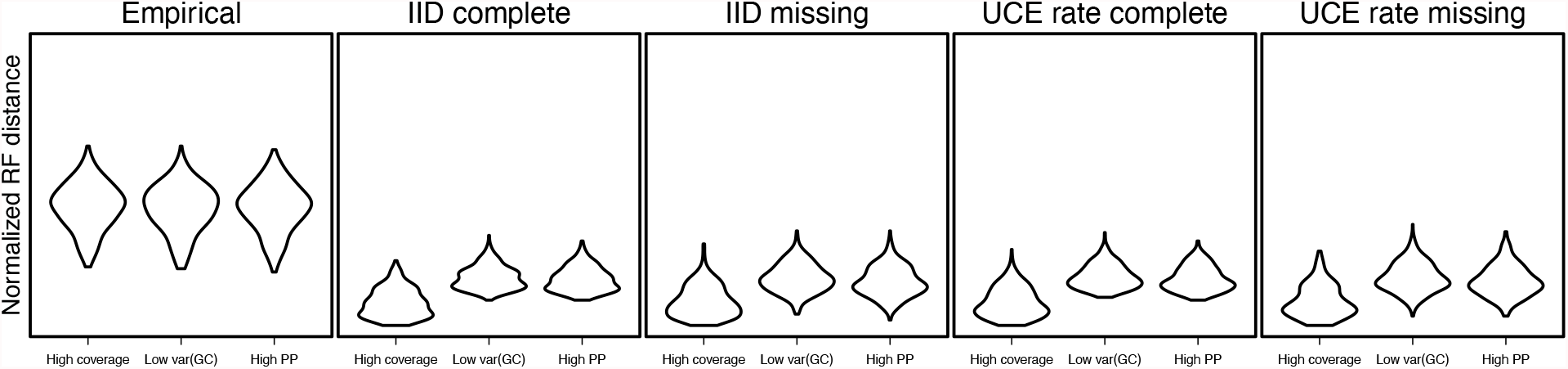
Gene tree discordance measured using the normalized Robinson-Foulds (RF) distance between several reference trees and the single gene trees. As reference trees, we used the the maximum a posterior (MAP) phylogeny using the three different data subsets (high coverage, low variance in GC content, and high posterior probability of *Photinus+Ellychnia* being monophyletic). The left panel shows the frequency of the RF-distance for the empirical dataset of Martin et al. (2019). The second panel shows the RF-distance for the simulated dataset with complete sequences and homogeneous (*i*.*e*., independent and identically distributed, IID) highly variables sites. The third panel shows the RF-distance for the simulated dataset with missing sequences and homogeneous highly variable sites. The fourth panel shows the RF-distance for the simulated dataset with complete sequences and systematically distributed (*i*.*e*., akin to the empirical AHE dataset) highly variables sites. The right panel shows the RF-distance for the simulated dataset with missing sequences and systematically distributed highly variable sites. Neither of our simulation conditions are a strongly negative impact on gene tree discordance.

We observed that missing data has an impact on the distribution of posterior probabilities of certain clades being monophyletic (Figures S13-S16). Some clades, for example, Phosphaeni, Luciolinae and *Curtos*, are always rejected if complete sequences were used in the simulations (Figures S13 and S15). When we\ used sequences with missing data in our simulations, and pruned sequences with less than 50% non-missing sites, then we obtain high posterior probabilities for some loci that these clades were indeed monophyletic (Figures S14 and S16). The latter results agree more with our empirical analysis.

Note that we used the phylogeny which we inferred for the concatenated alignment to simulate dataset (Figure 7). This phylogeny has some clades, for example *Curtos*, as monophyletic. Single loci were the possibly wrongly placed *Curtos sp* was removed because of missing sites resulted in high posterior probabilities of *Curtos* being monophyletic. The same influence of pruning taxa because of missing sites can be observed for other problematic clades. Thus, some taxa with high fractions of missing sites might be placed incorrectly because the sequences could be of lower quality and with higher errors.

**Figure 7:**
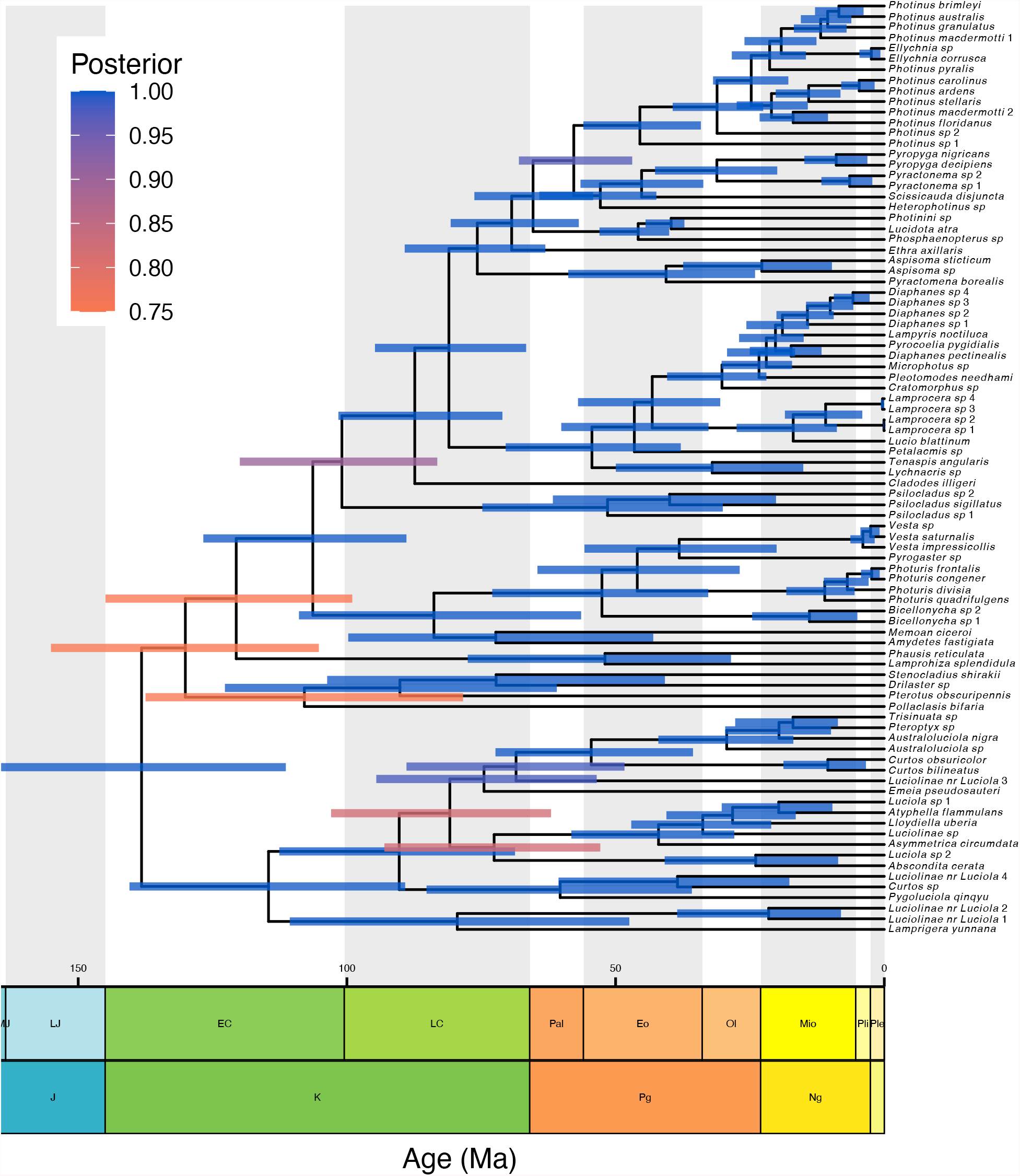
Time-calibrated phylogeny of Lampyridae. Estimated time-calibrated Lampyridae phylogeny under the fossilized birth-death-range process using the high posterior probability of *Photinus* being monophyletic data subset. The plot was generated using the R package `RevGadgets` (Tribble et al. 2022). The bars around the node show the 95% credible interval of the node ages. Bars are colored according to the posterior probability of the node.

### Time-Calibrated Lampyridae Phylogeny

#### Topology of Lampyridae

The estimated topology of the dated Lampyrid tree is largely congruent with that presented in Martin et al. (2019), which was generated with the same AHE dataset without fossil calibration. However, there are some minor differences between the two trees, distributed throughout the phylogeny (Figure 7, Supplementary Table S2). While the core topology of the dated tree supports the recent higher-level classification (Martin et al. 2019), these minor topological differences have the potential to influence evolutionary analyses, thus highlighting the need for expanded taxonomic and phylogenomic investigations of fireflies, as well as development of robust phylogenomic methods in general.

The support for the named clades using the three concatenated data subsets largely matches the support from the single gene tree analyses (Figure 2 and S17). The concatenated analyses inferred trees with extremely high support; the posterior probabilities of the named clades were either 0.0 or 1.0. This strong support could be inflated posterior probabilities instead of true signal. The selected four MCMC replicates show identical posterior probabilities, indicating convergence of the MCMC analyses.

The most interesting results are obtained for the clade that received ambiguous support from the single gene trees: *Photinus+Ellychnia, Australoluciola* and Ototretinae. It is expected that we received high posterior support for *Photinus+Ellychnia* being monophyletic for the data subset with the loci support *Photinus+Ellychnia* monophyly with at least 0.95 posterior probability. Reassuringly, the other two data subsets also recovered *Photinus+Ellychnia* to be monophyletic. It is therefore most probable that *Photinus+Ellychnia* is indeed monophyletic. In all our analysis we recovered both *Ellychnia* itself being monophyletic and *Photinus* + *Ellychnia* being monophyletic, indicating that *Photinus* is paraphyletic (Figure 7). The inclusion of *Ellychnia* within *Photinus* has been reported previously (Stanger-Hall et al. 2007; Stanger-Hall and Lloyd 2015; Martin et al. 2019) and has led Zaragoza-Caballero et al. (2020) to change *Ellychnia* to *Photinus*.

All three data subsets agreed that *Australoluciola* is not monophyletic. In the AHE dataset from Martin et al. (2019) there are only two species belonging to *Australoluciola* (see Table S1). Our inferred results show *Australoluciola* being paraphyletic with *Pteroptyx sp* and *Trisinuata sp* nested within, agreeing with previous results by Jusoh et al. (2018). Ototretinae also consisted of only two species (*Drilaster sp* and *Stenocladius shirakii*) in the AHE dataset by Martin et al. (2019). Ototretinae was inferred to be monophyletic using the *Photinus+Ellychnia* 0.95 posterior probability data subset, but was not found to be monophyletic using the other two data subsets (Figure S17).

#### Divergence Times of Lampyridae

We inferred a time-calibrated phylogeny of Lampyridae. Our estimate of the crown age of Lampyridae is 139.85 Ma with a 95% credible of [108.43, 165.68]. Our estimated crown age is older than most previous estimates (Table 2). Toussaint et al. (2017) showed that previous divergence time estimates of McKenna et al. (2015) are likely underestimates. Specifically, McKenna et al. (2015) estimated a crown age of Elateroidea of 166.18 (151.83–181.57) while Toussaint et al. (2017) estimated a crown age of 246.02 (231.35–260.12).

**Table 2:**
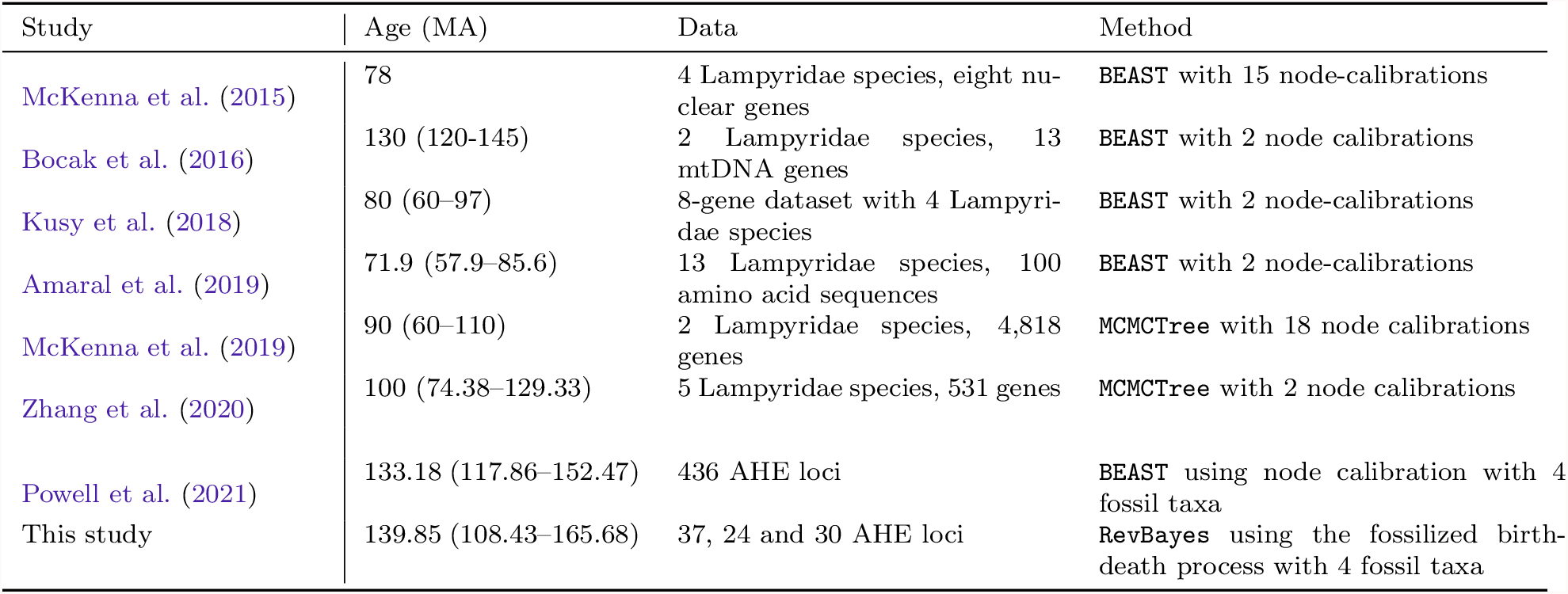
Fossils and calibration constraints for the time-calibrated divergence time analysis.

| Study | Age (MA) | Data | Method |
| --- | --- | --- | --- |
| McKenna et al. (2015) | 78 | 4 Lampyridae species, eight nuclear genes | BEAST with 15 node-calibrations |
| Bocak et al. (2016) | 130 (120-145) | 2 Lampyridae species, 13 mtDNA genes | BEAST with 2 node calibrations |
| Kusy et al. (2018) | 80 (60–97) | 8-gene dataset with 4 Lampyridae species | BEAST with 2 node-calibrations |
| Amaral et al. (2019) | 71.9 (57.9–85.6) | 13 Lampyridae species, 100 amino acid sequences | BEAST with 2 node-calibrations |
| McKenna et al. (2019) | 90 (60–110) | 2 Lampyridae species, 4,818 genes | MCMCTree with 18 node calibrations |
| Zhang et al. (2020) | 100 (74.38–129.33) | 5 Lampyridae species, 531 genes | MCMCTree with 2 node calibrations |
| Powell et al. (2021) | 133.18 (117.86–152.47) | 436 AHE loci | BEAST using node calibration with 4 fossil taxa |
| This study | 139.85 (108.43–165.68) | 37, 24 and 30 AHE loci | RevBayes using the fossilized birth-death process with 4 fossil taxa |

Most previous analyses used only very few Lampyridae species (up to five species) which could possibly bias crown age estimates if the true crown group was not sampled. Furthermore, most previous studies should not be considered as independent evidence, as for example Zhang et al. (2020) uses divergence times for calibrations which were estimated by Zhang et al. (2018).

Our divergences times overlap with the recent estimates by Powell et al. (2021) using the same AHE dataset. The main difference in the two analysis are (i) the number of loci used (436 in Powell et al. (2021) and up to 37 loci in this study), (ii) the chosen fossil calibrations, (iii) the fossil calibration approach (node dating vs the fossilized-birth-death process), and (iv) the software `BEAST` vs `RevBayes`. It is very reassuring to see the large overlap in divergence time estimates despite these differences in the performed analysis.

Our divergence times are robust for the majority of clades when comparing the three different data subsets (Figure S18). If the clades were identified as being monophyletic for all three data subsets (Figure S17), then also the estimated crown ages were identical (Figure S18). However, when the clades were not found to be monophyletic, then the clade ages also differed (*e*.*g*., Ototretinae). This result is not surprising as the crown age is defined as the most recent common ancestor for the selected taxa, and if this most recent common ancestor includes different species then the interpretation of this ancestor and its age should be different.

Our sensitivity analysis of including and excluding the recently published *Photinus* fossil, †*Photinus kazantsevi* (Alekseev 2019), yielded largely identical results (Figure 8). †*Photinus kazantsevi* was dated to be 33.9 to 37.2 million years old. Our estimated crown age of *Photinus* was between 32 and 63 Ma, including and excluding †*Photinus kazantsevi*. This result gives us confidence about our estimate of the crown age of *Photinus*.

**Figure 8:**
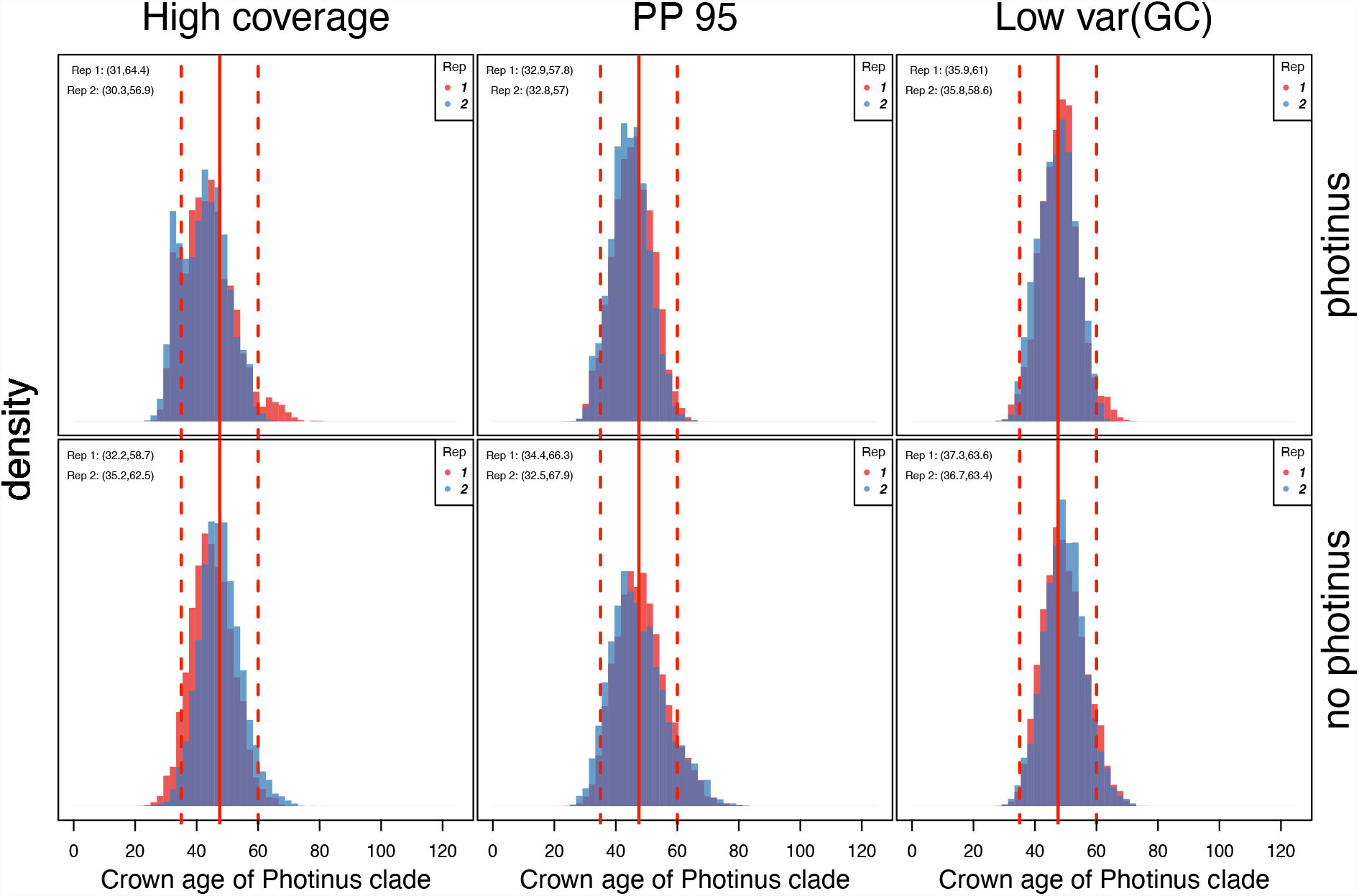
Estimated crown age of *Photinus*. We show the crown age of the *Photinus* clade for the three data subsets with (top row) and without (bottom row) using the †*Photinus kazantsevi*. First, we observe that the data subset has little impact on the estimated *Photinus* crown age. Second, the usage of †*Photinus kazantsevi* does not significantly impact the *Photinus* crown age estimate, which corroborates the †*Photinus kazantsevi* placement and *Photinus* crown age.

## Discussion

### Divergence time estimation using genomic data

The objective of this study was to estimate a time-calibrated phylogeny of fireflies using genomic data. The computational demand was extremely high prohibiting the full use of the 436 loci combined with adequate full-likelihood Bayesian divergence time estimation methods (*e*.*g*., adequately partitioned substitution model, relaxed clock model and fossilized birth-death process). Even when we used a smaller subset of the data, *i*.*e*., 24 to 37 loci, the computational demand was still extremely high (several weeks to months for a single analysis) but manageable. Until faster methods are available, we can only resort to using a subset of data if we wish to use full-likelihood methods for divergence time estimation. However, until now, there are no clear guidelines on how to select the best data subset for divergence time estimation. Here, we constructed three data subsets and explored several characteristics of the data. Nevertheless, understanding the lack of a clear correlation of our data subsets to phylogenetic accuracy, has proven to be challenging.

First, we observed that our inferred phylogenies from the three data subsets are mostly identical. That means that different data subset can converge on the same phylogeny and this could indicate support that the inferred phylogeny is robust. Other studies had shown that different data sources (*e*.*g*., exons, ultra conserved elements and transcriptomic sequences) yield different phylogenies, *e*.*g*., Betancur-R et al. (2019). In our study with used the same data source but different data subset. Thus, the difference in results of phylogenomic studies could originate rather from the data source than the amount of data.

Second, divergence time estimates seem robust to the chosen data subset if the inferred topology agreed. It is not surprising that a clade, which was inferred to be monophyletic for one data subset but not for another data subset, obtained a different crown age estimate (*e*.*g*., Ototretinae). Therefore, we conclude that it is more important to focus first on robust estimation of phylogeny using different data subsets or methods. Once we understand how to select data subsets to produce reliable phylogenies, then we can safely use the same data subsets for divergence time estimation. Our study raises some important aspects where we need to improve our inference of phylogeny.

### Unrealistic gene tree discordance

Our analyses of the 436 AHE loci revealed strong gene tree discordance. Such large amounts of gene tree discordance are not expected even when allowing for incomplete lineage sorting. Richards et al. (2018) showed that discordance between mitochondrial loci and nuclear loci is equally large, suggesting that much of the apparent gene tree discordance originates from methodical factors and biological factors. Our simulation study corroborates these findings: we cannot explain the observed gene tree discordance with incomplete lineage sorting or phylogenetic uncertainty.

In our analyses, we did not clean the AHE dataset but took the data as given. There are several possible sources of error which we did not check. For example, we did not assess orthology and did not check for alignment errors. However, the published AHE dataset (Martin et al. 2019) was curated following current best practices and therefore should reflect the state of the field.

In the last decade, we have seen several debates about using concatenation or coalescent-based species tree approaches (for a recent review see Bravo et al. 2019). Our observed higher accuracy of the concatenated analyses over the single gene trees is surprising and provides empirical evidence against standard theoretical expectations. We expect that there is recombination between loci and therefore that concatenation approaches can be misleading (Degnan and Rosenberg 2009). Instead, we find that the information within single loci is misleading and, when concatenated, the noise is canceled out to leave an improved phylogenetic signal (*e*.*g*.,, recovery of *Photinus + Ellychnia* as monophyletic). We do not advocate that concatenation approaches are philosophically or theoretically superior, but instead we notice that current single gene tree estimation methods are plagued by high gene tree error (Bossert et al. 2021) and thus we find a strong empirical violation of the assumptions of the multispecies coalescent process Reid et al. (2014). We need to understand and alleviate the gene tree error before we can continue with the debate on coalescent-based species tree approaches. Finally, even if both concatenation and summary-based multispecies coalescent approach show robustness to noise in single gene tree estimates (Molloy and Warnow 2018), noise in single gene tree estimates are highly problematic when used for ancient population size estimation and inference of gene flow (Kutschera et al. 2014). Our simulations show that it is possible to infer gene trees correctly and using a concatenation approach hides the underlying problems in gene tree inference.

### Filtering loci

Previous studies have shown that loci can be filtered to increase phylogenetic accuracy (*e*.*g*., Alda et al. 2019). However, previous studies mainly focused on the resulting species tree and not on single genes (*e*.*g*., Leite et al. 2021). Overall, we could not identify a single summary statistics as a reliable predictor for phylogenetic accuracy (Figure 3). Thus, we could not use single summary statistics as data filtering criteria for robust phylogenetic inference. It is possible that some phylogenetic relationships of Lampyridae require revision and thus our proxies of phylogenetic accuracy are misleading. Clearly more work is needed if these summary statistics of the data are used for selecting data subsets. Our results show some promise and single summary statistics could be used to detect outliers which are then removed to clean the dataset (Figure 3). Additionally, a combination of summary statistics might provide a fruitful future approach.

### Posterior predictive simulations

Posterior predictive distributions to test model adequacy have not been routinely applied in phylogenomic studies (Brown and Thomson 2018). If a model is not adequate for a given dataset, then we cannot guarantee the accuracy and robustness of the estimates. With genomic data, our hope is that for some loci we have adequate phylogenetic substitution models while for other loci we do not. Such a result would allow us to proceed with the subset of loci that we can model adequately. Unfortunately, our results show that we do not have adequate phylogenetic substitution models for any of the 436 AHE loci. This fact should not be downplayed and we direly need more accurate substitution models. First, we need to develop a better understanding of posterior predictive distribution and the expected behavior under simulations. Second, we need more and better summary statistics that will guide us in the development of more accurate phylogenetic substitution models. Third, we need to modify our standard phylogenetic pipelines to include more complex substitution models, for example, within-locus partitioning (Freitas et al. 2021), site-heterogeneous substitution models (Lartillot et al. 2007; Wu et al. 2013) and Markov modulate Markov model (Baele et al. 2021). Then, we should revisit both the gene tree accuracy as well as gene tree model adequacy. Until then, we cannot say with confidence why we observe so much gene tree discordance, whether biological or methodological.

#### Missing data and summary statistics

We investigated here if missing sequence data is a problem for phylogenetic inference. In theory, missing sequence data does not pose a problem if sufficient information is retained (Philippe et al. 2004; Lemmon et al. 2009). Imagine that we would add another column to our data matrix but this column consists only of missing sites. In that case, we have not added any information to our data and in fact the likelihood score remains the same after adding this column of missing data (Felsenstein 2004). Hence, missing data in theory is not problematic.

Previous studies have investigated the impact of missing data using simulation studies (Philippe et al. 2004; Wiens 2003; Lemmon et al. 2009). A challenge to evaluate missing data is how the missing data is distributed. If we randomly place missing data in the data matrix, then this has only a minor impact on our ability to correctly infer the true phylogeny. Here we used an empirically informed approach and removed sites using the same positions as in the original data matrix (see Supplementary Figures S2-S4). In our simulations it occurred that complete sequences were missing and certain regions (the boundary of the sequences) have higher prevalence of missing data. This uneven distribution of missing data has the effect that some taxa cannot be placed accurately and the entire gene tree is erroneous. Therefore, it was necessary to remove taxa for a locus with too few non-missing sites, *e*.*g*., we removed taxa with fewer than 50% sites.

Our simulations and results confirm theoretical expectations that missing sites do not bias phylogenetic inference (Figure 6). However, we observed that missing sites do bias distributions of summary statistics (Figure 4) and a non-uniform distribution of highly variable sites together with a non-uniform distribution of missing sites can bias posterior predictive distributions (Figure 5). This study was the first study to explore missing data for posterior predictive distributions. In most cases, summary statistics are not explicitly defined for missing data. For example, how should the GC content be computed for a sequence where 50% of the sites are missing? We resolved the issue by computing the GC content of only the non-missing sites, although missing sites could be more concentrated at GC rich regions (Beauclair et al. 2019). Similarly, how should the minimum distance between two sequence which do not overlap be computed? Our approach was to simply omit these sequences. These two examples show the importance of simulating missing data with the same distribution as the observed data to not bias summary statistics. Since the prevalence of missing data increases for phylogenomic studies, we need to find better solutions to incorporate missing sequence data into our analyses and summary statistics, both for filtering as well as model adequacy testing.

## Conclusions

The primary aim of this study was to estimate a time-calibrated phylogeny of Lampyridae. We used the previously published 436 AHE loci from Martin et al. (2019). To calibrate the phylogeny, we employed the recently developed fossilized birth-death-range process (Stadler et al. 2018) together with standard relaxed-clock models (Drummond et al. 2006) in a Bayesian framework, as implemented in the software `RevBayes` (Höhna et al. 2016). Full Bayesian relaxed-clock divergence time estimation analyses cannot handle datasets with hundreds of loci without sacrificing model complexity. Instead, we selected three different data subsets and found that divergence time estimates agreed for all clades that were identical between analyses (Figure S18). We estimated a crown age of Lampyridae of 139.85 [108.43, 165.68] Ma which is considerably older than some previous estimates (Kusy et al. 2018; Zhang et al. 2018; Amaral et al. 2019; McKenna et al. 2019) but matches recent findings of earliest fossils belonging to Lampyridae (Kazantsev 2015) and is in agreement with some other studies (Bocak et al. 2016; Toussaint et al. 2017; Zhang et al. 2020; Powell et al. 2021) obtained from taxonomically broader studies. Thus, divergence time estimation using hundreds of loci is robust if a representative data subset is chosen. Previous results on topological disagreement depending on data filtering (*e*.*g*., Kuang et al. 2018; McLean et al. 2019) apply to divergence time estimation too.

In the process of selecting robust data subsets, we investigated the phylogenetic accuracy of single AHE loci. We found an unexpected amount of gene tree discordance (Figure 6). We explored the impact of incomplete lineage sorting, missing sequence data and systematic distribution of highly variable sites using simulations. The observed gene tree discordance cannot be explained due to incomplete lineage sorting. Instead, the gene tree discordance most likely originates from data errors (*e*.*g*., paralogs and poor alignments) or model inadequacy (Figure 5). Surprisingly, our standard phylogenetic substitution models are not adequate for even a single AHE locus. We showed that this model inadequacy is not due to missing data (Figure 5) although missing data influence the distribution of summary statistics (Figure 4). More work on understanding the causes of the apparent gene tree discordance is needed. It is paramount to have robust gene trees not only for phylogeny and divergence time estimation but also to draw any conclusions about biological processes such as incomplete lineage sorting, horizontal gene transfer and gene flow.

## Acknowledgements

This research was supported by the Deutsche Forschungsgemeinschaft (DFG) Emmy Noether-Program HO 6201/1-1 awarded to SH.

