## Supplementary Material for "A Time-calibrated Firefly (Coleoptera: Lampyridae) Phylogeny: Using Genomic Data for Divergence Time Estimation"

### A Time-calibrated Firefly (Coleoptera: Lampyridae) Phylogeny: Using Ultra Conserved Elements for Divergence Time Estimation

#### Supplementary Material

R.H. Time-calibrated Firefly Phylogeny

Sebastian Höhna[1,2], Sarah E. Lower[3], Pablo Duchen[4], and Ana Catalán[1,5]

[1] *GeoBio-Center, Ludwig-Maximilians-Universität München, 80333 Munich, Germany*

[2] *Department of Earth and Environmental Sciences, Paleontology & Geobiology, Ludwig-Maximilians-Universitt Mnchen, 80333Munich, Germany*

[3] *Department of Biology, Bucknell University, Lewisburg, PA 17837, U.S.A.*

[4] *Department of Computational Biology, University of Lausanne, 1015 Lausanne, Switzerland*

[5] *Division of Evolutionary Biology, Ludwig-Maximilians-Universität München Planegg-Martinsried 82152, Germany*

To whom correspondence should be addressed:

Sebastian Höhna
GeoBio-Center
Department of Earth and Environmental Science
Palaeontology & Geobiology
Ludwig-Maximilians-Universität München
Richard-Wagner-Straße 10
80333 München
Germany

Phone: +49 (0)89 / 2180-6615

### Genera, Tribes and Sub-families

**Table S1: Genera, Tribes and Sub-families used to assess gene-tree error and clade ages.**

| Name | Species |
|---|---|
| Psilocladus | Psilocladus sigillatus, Psilocladus sp 1, Psilocladus sp 2 |
| Vesta | Vesta impressicollis, Vesta saturnalis, Vesta sp |
| Amydetinae | Amydetes fastigiata, Cladodes illigeri, Ethra axillaris, Memoan ciceroi, Psilocladus sigillatus, Psilocladus sp 1, Psilocladus sp 2, Vesta impressicollis, Vesta saturnalis, Vesta sp |
| Lamprohizinae | Lamprohiza splendidula, Phausis reticulata |
| Aspisoma | Aspisoma sp, Aspisoma sticticum |
| Pyractomena | Pyractomena borealis, Pyractonema sp 1, Pyractonema sp 2 |
| Cratomorphini | Aspisoma sp, Aspisoma sticticum, Cratomorphus sp, Pyractomena borealis, Pyractonema sp 1, Pyractonema sp 2 |
| Lamprocera | Lamprocera sp 1, Lamprocera sp 2, Lamprocera sp 3, Lamprocera sp 4 |
| Lamprocerini | Lamprocera sp 1, Lamprocera sp 2, Lamprocera sp 3, Lamprocera sp 4, Lucio blattinum, Lychnacris sp, Tenaspis angularis |
| Diaphanes | Diaphanes pectinealis, Diaphanes sp 1, Diaphanes sp 2, Diaphanes sp 3, Diaphanes sp 4 |
| Lampyrini | Diaphanes pectinealis, Diaphanes sp 1, Diaphanes sp 2, Diaphanes sp 3, Diaphanes sp 4, Lampyris noctiluca, Microphotus sp, Petalacmis sp, Pleotomodes needhami, Pyrocoelia pygidialis |
| Ellychnia | Ellychnia corrusca, Ellychnia sp |
| Photinus | Photinus sp 1, Photinus sp 2, Photinus floridanus, Photinus macdermotti 1, Photinus stellaris, Photinus ardens, Photinus carolinus, Photinus pyralis, Ellychnia sp, Ellychnia corrusca, Photinus macdermotti 2, Photinus granulatus, Photinus australis, Photinus brimleyi |
| Pyropyga | Pyropyga decipiens, Pyropyga nigricans |
| Photinini | Photinini sp , Ellychnia corrusca, Ellychnia sp, Heterophotinus sp, Lamprigera yunnana, Photinus sp 1, Photinus sp 2, Photinus floridanus, Photinus macdermotti 2, Photinus stellaris, Photinus ardens, Photinus carolinus, Photinus pyralis, Ellychnia sp, Ellychnia corrusca, Photinus macdermotti 1, Photinus granulatus, Photinus australis, Photinus brimleyi, Pyropyga decipiens, Pyropyga nigricans |
| Phosphaenini | Lucidota atra, Phosphaenopterus sp |
| Lampyrinae | Aspisoma sp, Aspisoma sticticum, Cratomorphus sp, Pyractomena borealis, Pyractonema sp 1, Pyractonema sp 2, Lamprocera sp 1, Lamprocera sp 2, Lamprocera sp 3, Lamprocera sp 4, Lucio blattinum, Lychnacris sp, Tenaspis angularis, Diaphanes pectinealis, Diaphanes sp 1, Diaphanes sp 2, Diaphanes sp 3, Diaphanes sp 4, Lampyris noctiluca, Microphotus sp, Petalacmis sp, Pleotomodes needhami, Pyrocoelia pygidialis, Photinini sp , Ellychnia corrusca, Ellychnia sp, Heterophotinus sp, Lamprigera yunnana, Photinus sp 1, Photinus sp 2, Photinus floridanus, Photinus macdermotti 2, Photinus stellaris, Photinus ardens, Photinus carolinus, Photinus pyralis, Ellychnia sp, Ellychnia corrusca, Photinus macdermotti 1, Photinus granulatus, Photinus australis, Photinus brimleyi, Pyropyga decipiens, Pyropyga nigricans, Lucidota atra, Phosphaenopterus sp |
| Australoluciola | Australoluciola nigra, Australoluciola sp |
| Curtos | Curtos sp, Curtos bilineatus, Curtos obsuricolor |
| Luciola | Luciola sp 1, Luciola sp 2 |
| Luciolini | Abscondita cerata, Asymmetrica circumdata, Atyphella flammulans, Australoluciola sp, Australoluciola nigra, Curtos sp, Curtos bilineatus, Curtos obsuricolor, Emeia pseudosauteri, Lloydiella uberia, Luciola sp 1, Luciola sp 2, Pteroptyx sp, Pygoluciola qinqyu, Trisinuata sp |
| Luciolinae | Luciolinae sp, Luciolinae nr Luciola 1, Luciolinae nr Luciola 2, Luciolinae nr Luciola 3, Luciolinae nr Luciola 4, Abscondita cerata, Asymmetrica circumdata, Atyphella flammulans, Australoluciola sp, Australoluciola nigra, Curtos sp, Curtos bilineatus, Curtos obsuricolor, Emeia pseudosauteri, Lloydiella uberia, Luciola sp 1, Luciola sp 2, Pteroptyx sp, Pygoluciola qinqyu, Trisinuata sp |
| Ototretinae | Drilaster sp, Stenocladius shirakii |
| Bicellonycha | Bicellonycha sp 1, Bicellonycha sp 2 |
| Photuris | Photuris congener, Photuris divisia, Photuris frontalis, Photuris quadrifulgens |
| Photurinae | Bicellonycha sp 1, Bicellonycha sp 2, Photuris congener, Photuris divisia, Photuris frontalis, Photuris quadrifulgens, Pyrogaster sp |

### Sequence coverage

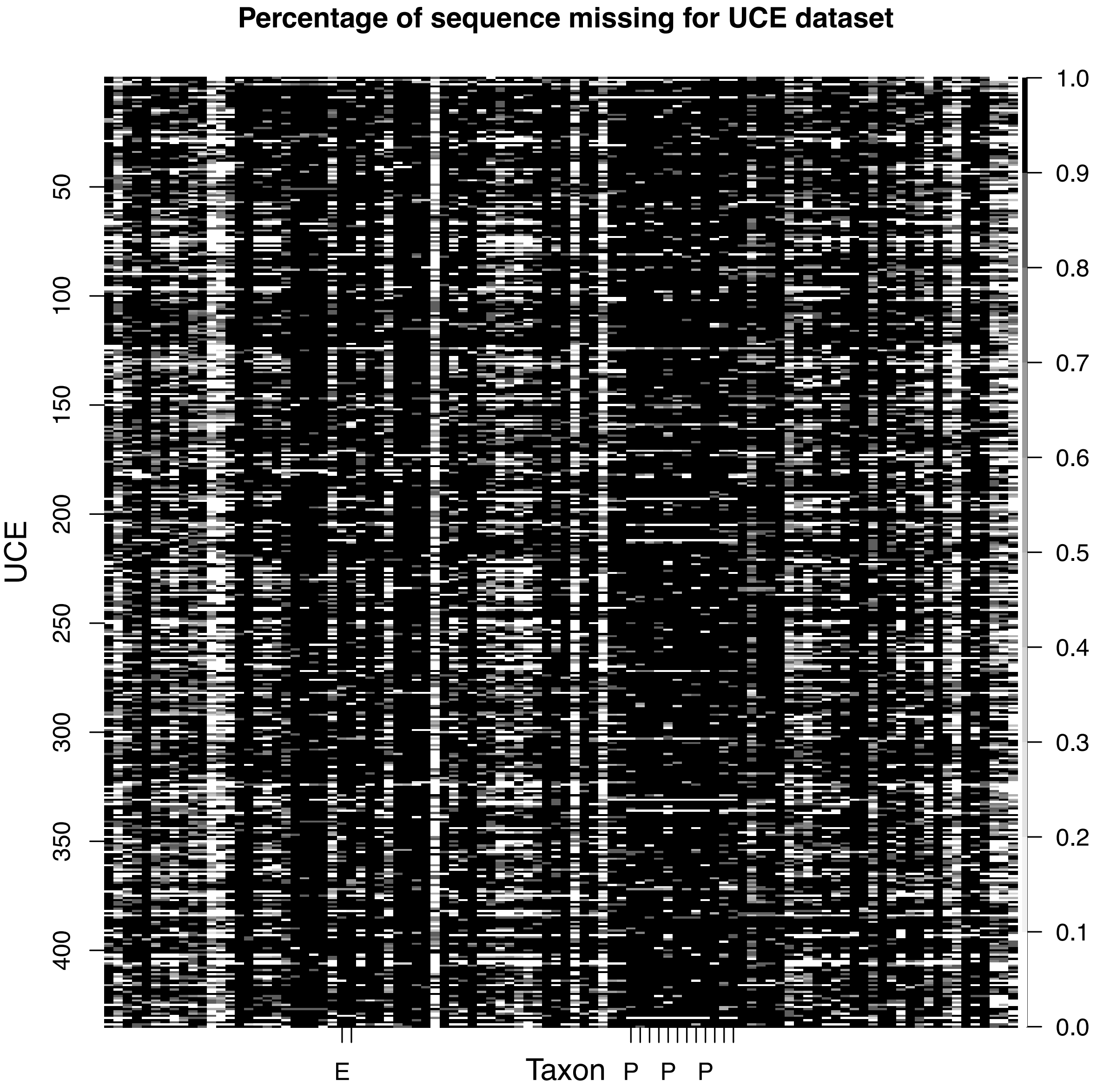


**Figure S1: Completeness of the AHE dataset of Martin et al. (2019).** We computed the percentage of sites missing per sequence. Black cells depict complete sequences and white cells depict entirely missing sequence. The gray shades depict the percentage in between. Each row represents one of the 436 AHE loci and each column represent one taxon. We highlighted the *Photinus* (P) and *Ellychnia* (E) species using tick marks.

3 # Simulated alignments

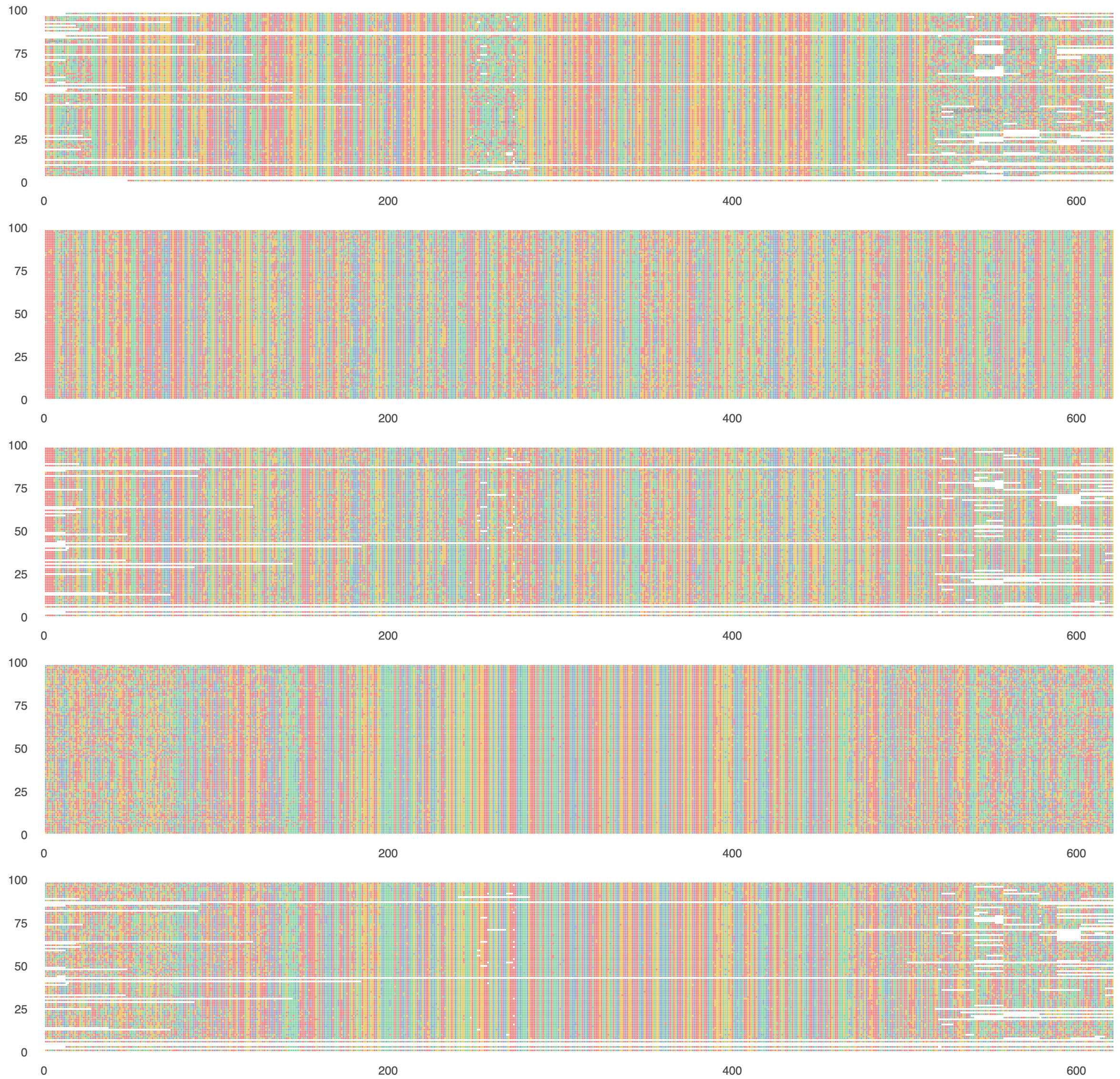


**Figure S2: First example locus for the simulations.** The top alignment shows the empirical data. The second row shows the simulated dataset with complete sequences and homogeneous (*i.e.,* independent and identically distributed, IID) highly variables sites. The third row shows the simulated dataset with missing sequences and homogeneous highly variable sites. The fourth row shows the simulated dataset with complete sequences and systematically distributed (*i.e.,* akin to the empirical UCE dataset) highly variables sites. The fifth row shows the simulated dataset with missing sequences and systematically distributed highly variable sites.

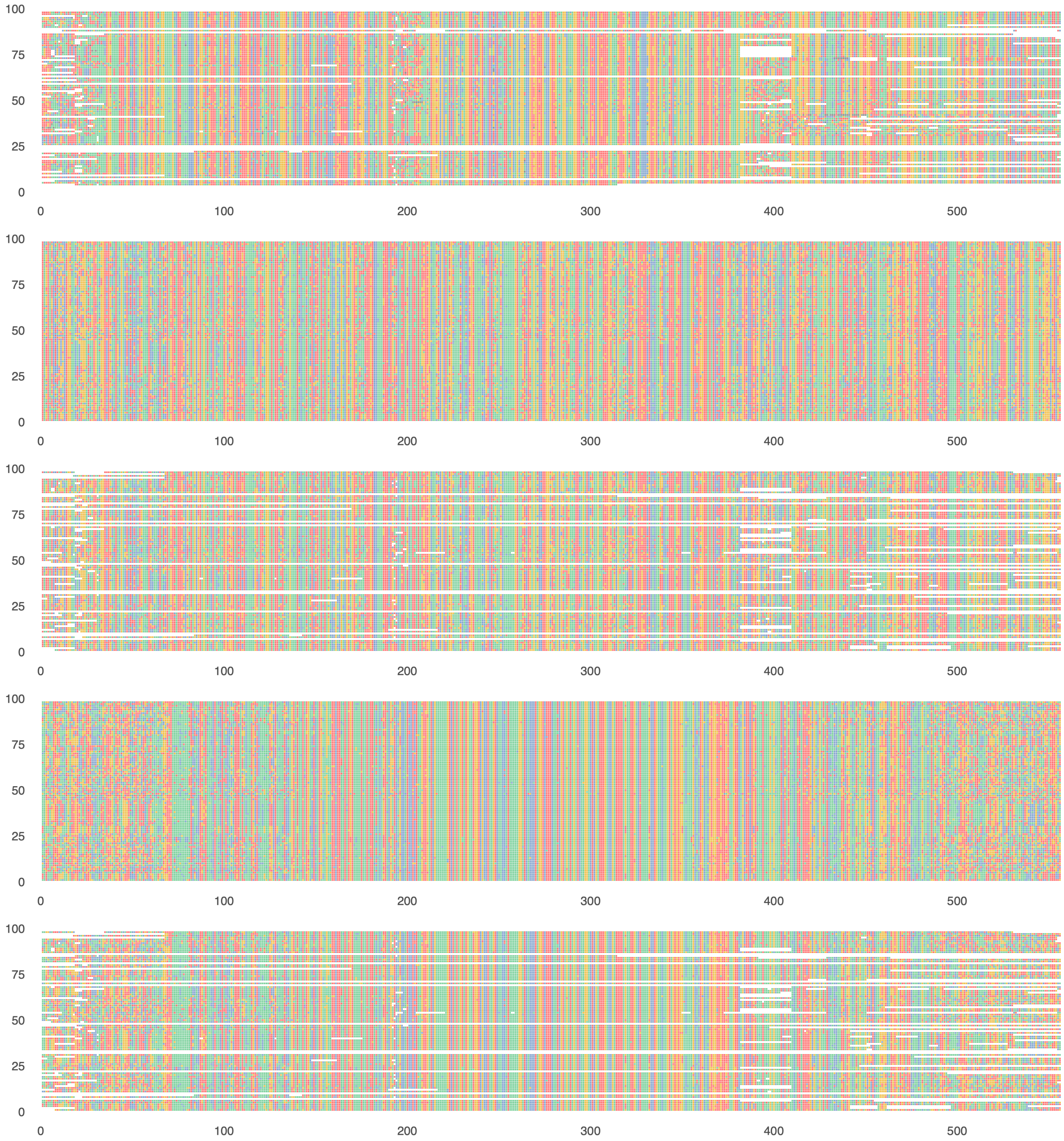


**Figure S3: Second example locus for the simulations.** The top alignment shows the empirical data. The second row shows the simulated dataset with complete sequences and homogeneous (*i.e.,* independent and identically distributed, IID) highly variables sites. The third row shows the simulated dataset with missing sequences and homogeneous highly variable sites. The fourth row shows the simulated dataset with complete sequences and systematically distributed (*i.e.,* akin to the empirical UCE dataset) highly variables sites. The fifth row shows the simulated dataset with missing sequences and systematically distributed highly variable sites.

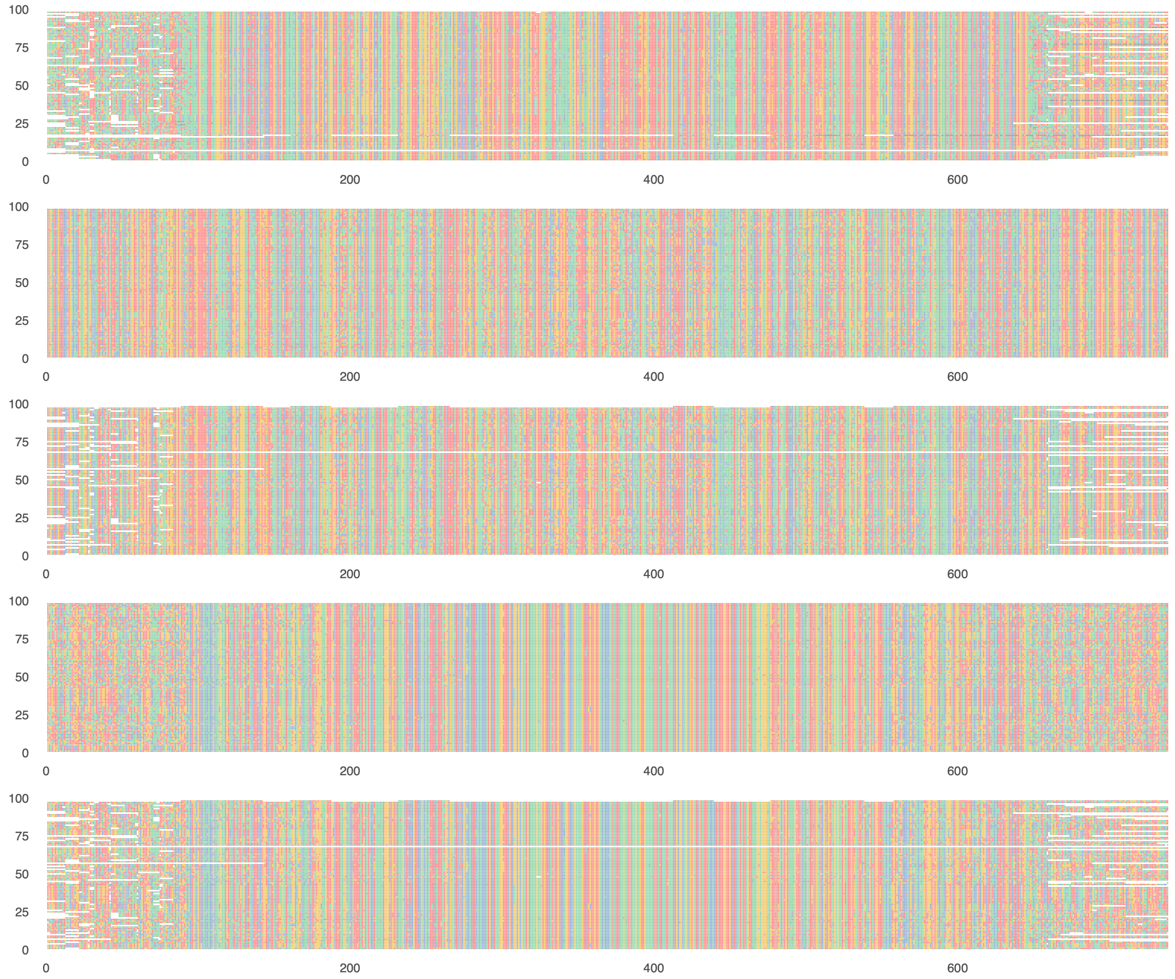


**Figure S4: Third example locus for the simulations.** The top alignment shows the empirical data. The second row shows the simulated dataset with complete sequences and homogeneous (*i.e.,* independent and identically distributed, IID) highly variables sites. The third row shows the simulated dataset with missing sequences and homogeneous highly variable sites. The fourth row shows the simulated dataset with complete sequences and systematically distributed (*i.e.,* akin to the empirical UCE dataset) highly variables sites. The fifth row shows the simulated dataset with missing sequences and systematically distributed highly variable sites.

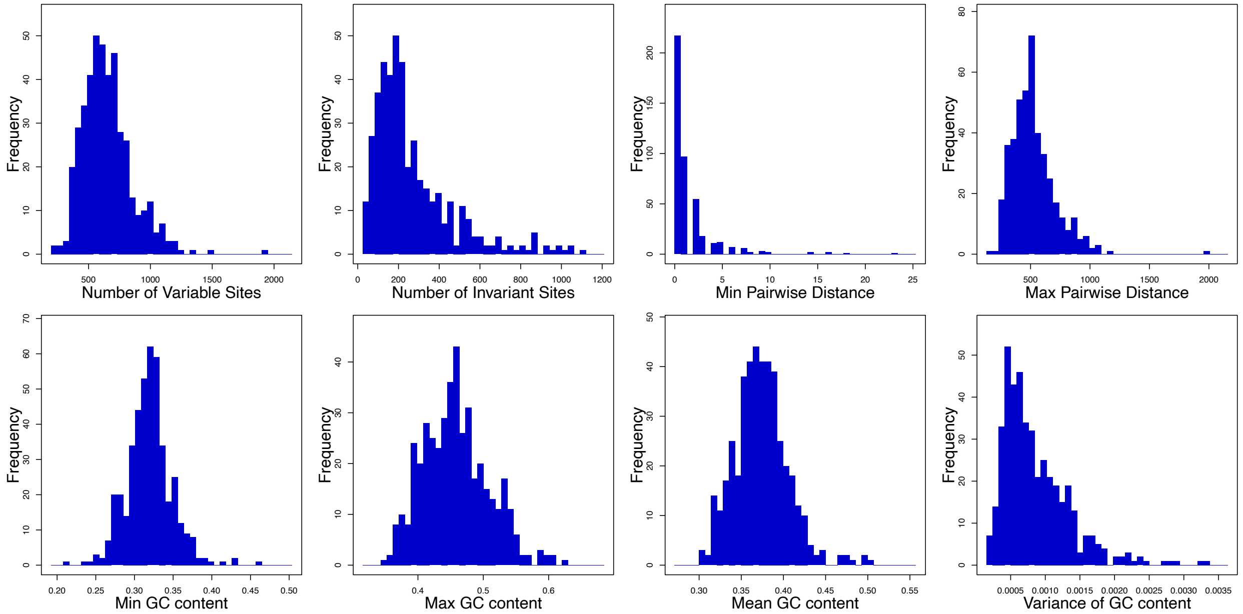


**Figure S5: Summary statistics obtained for the AHE dataset of Martin et al. (2019).** We computed the frequencies of summary statistics for the 436 AHE loci. Specifically, we compute the number of variable sites, number of invariant sites, minimum pairwise distance, maximum pairwise distance, minimum GC content, maximum GC content, mean GC content and variance of GC content.

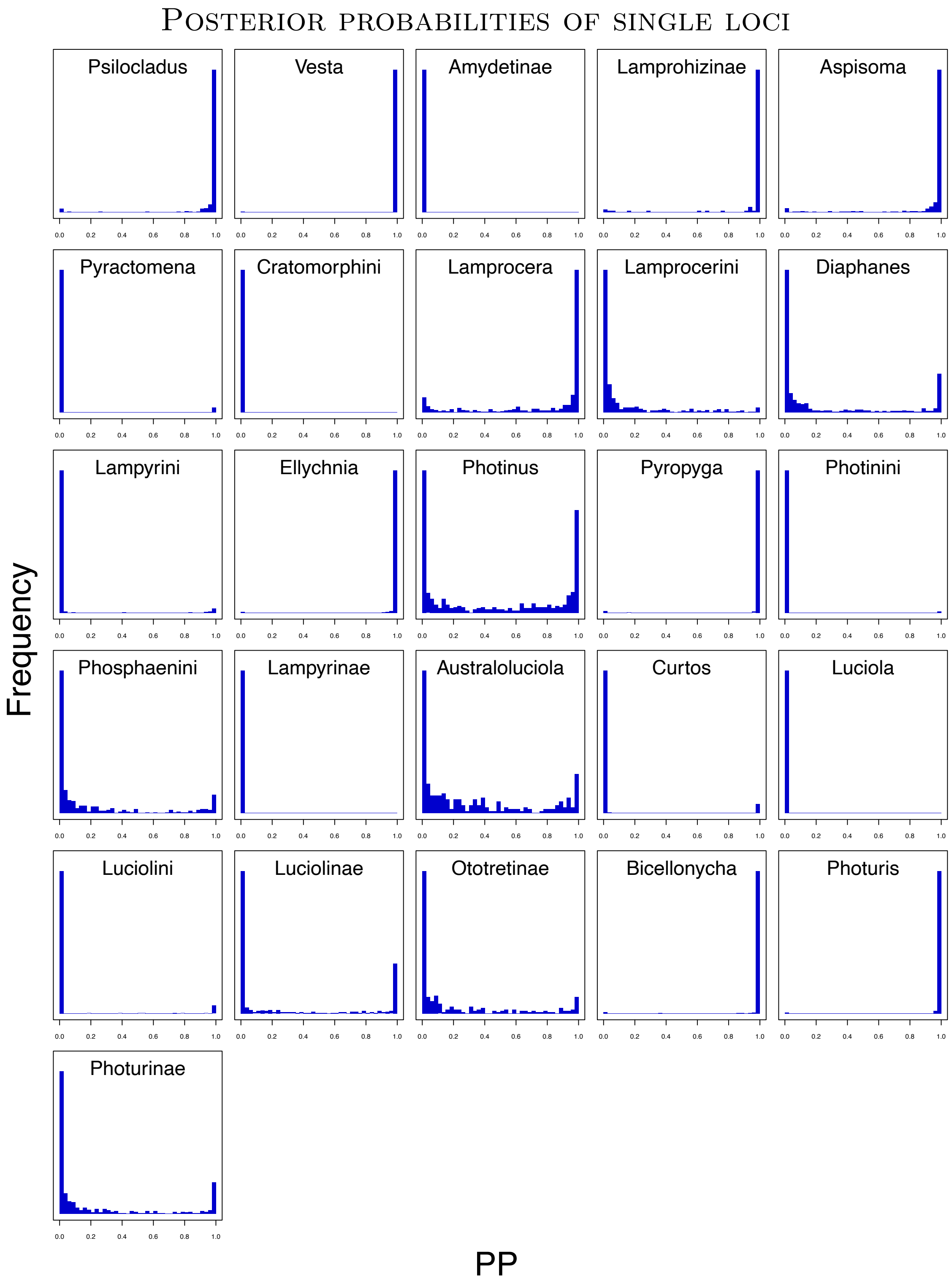


**Figure S6: Posterior probability of all 26 subfamilies, tribes and genera being monophyletic per AHE locus.** For each AHE locus, we computed the posterior probability that the clade is monophyletic. A histogram with most loci having a high posterior probability (*e.g., Vesta*) depicts strong support by the majority of AHE loci. Conversely, a histogram with most loci having a low posterior probability (*e.g., Photinus*) depicts strong support against monophyly by the majority of AHE loci. Four of the six subfamilies were not monophyletic. Only Lamprohizinae was found to be monophyletic for the majority of loci and Ototretinae was ambiguously supported. None of the six tribes was found to be monophyletic. Several genera are strongly rejected (*Pyractomena*, *Diaphanes*, *Photinus*, *Luciola* and *Curtos*).

### Correlation between sequence coverage and different clades being monophyletic

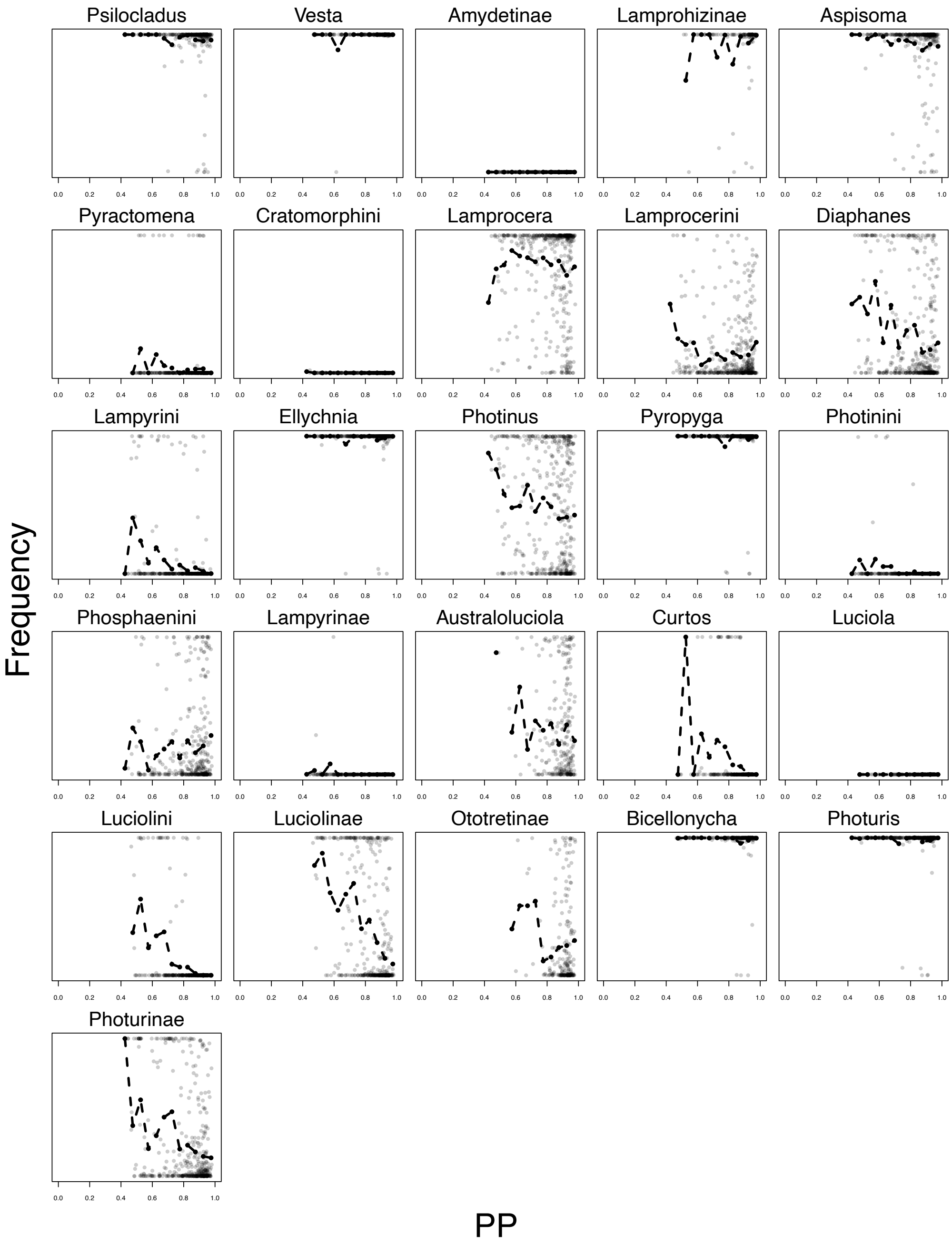


**Figure S7: Posterior probability of a clade being monophyletic per AHE locus versus the percentage of missing sites.** For each UCE locus, we computed the posterior probability that a given clade is monophyletic. We then compared the posterior probability of the clade being monophyletic to the percentage of missing sites for the locus. Each gray dot represents a single locus. The loci are plotted as opaque to better show clusters. The black dots connected by the black dashed line shows the rolling average of 20 loci.

### Correlation between summary statistics of the data and posterior support

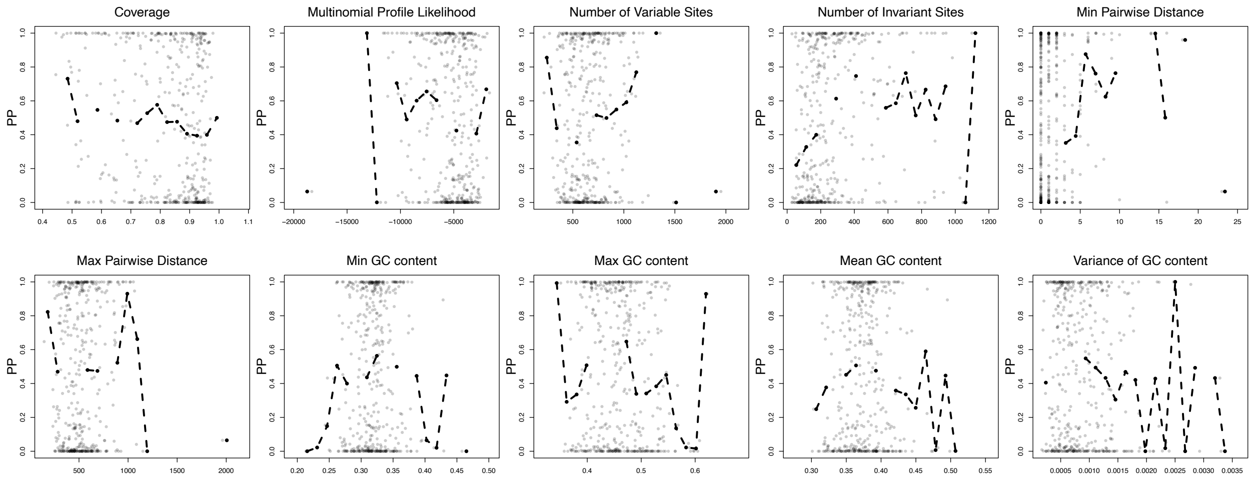


**Figure S8: Comparison of posterior probability of the clade *Photinus* being monophyletic to sequence coverage and other summary statistics obtained for the AHE dataset of Martin et al. (2019).** For each AHE locus, we computed the posterior probability that the clade *Photinus* is monophyletic. We computed the percentage of missing sites (coverage) as well as the summary statistics for the 436 AHE loci. Specifically, we compute the number of multinomial likelihood, variable sites, number of invariant sites, minimum pairwise distance, maximum pairwise distance, minimum GC content, maximum GC content, mean GC content and variance of GC content.

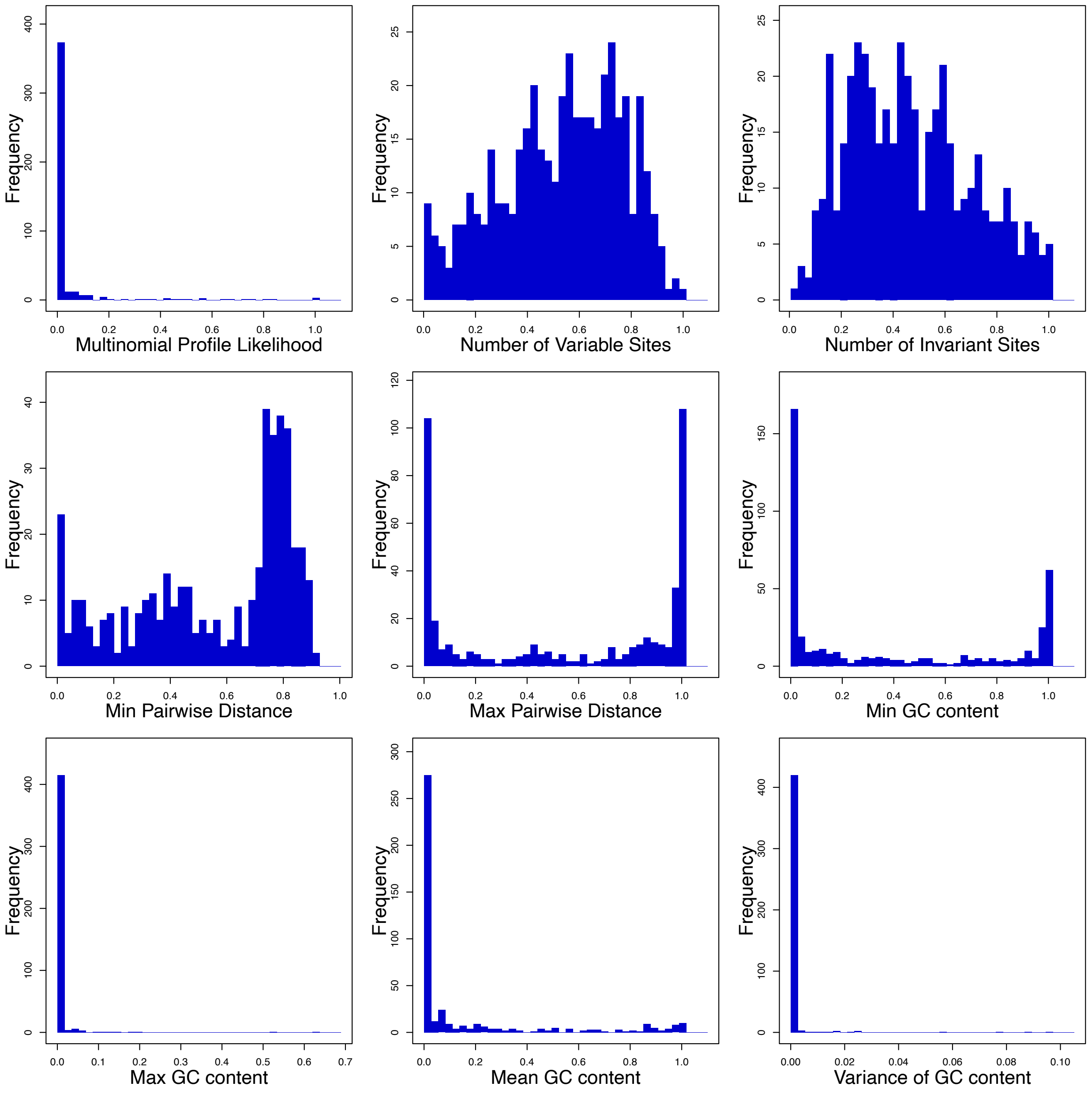


**Figure S9: Posterior predictive p-values for each UCE locus.** For each AHE locus, we simulated from the posterior predictive distribution and computed the posterior predictive p-value for various summary statistics. Specifically, we compute the number of variable sites, number of invariant sites, minimum pairwise distance, maximum pairwise distance, minimum GC content, maximum GC content, mean GC content and variance of GC content. Interestingly, most posterior predictive p-values show clear model violations, which could induce biased phylogenetic tree estimates. Surprisingly, even simple summary statistics that are usually modeled adequately, such as number of variable sites and number of invariant sites, are strongly violated.

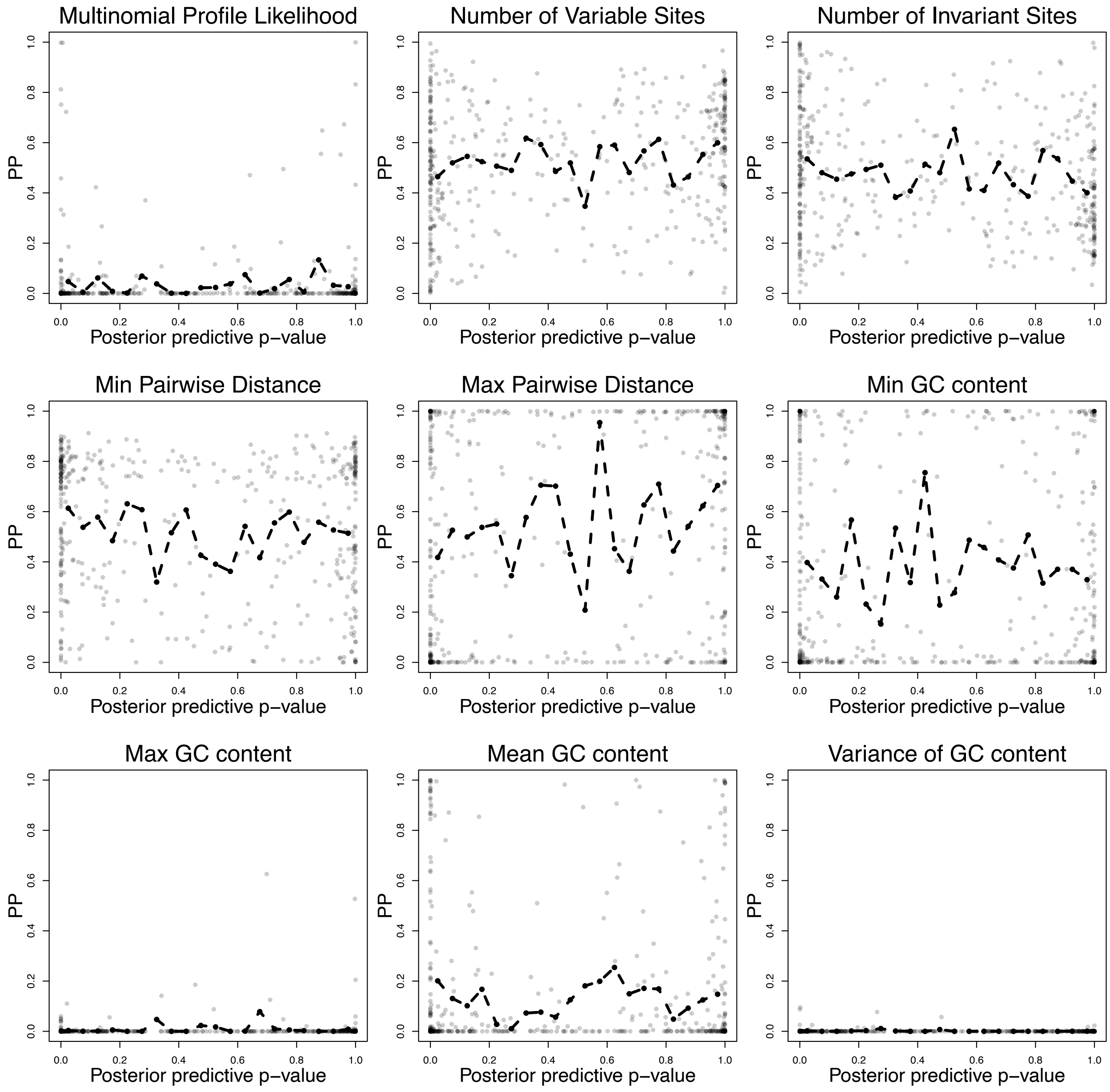


**Figure S10: Comparison of posterior probability of the clade *Photinus* being monophyletic to posterior predictive p-values.** The dashed black line shows a rolling average. There does not seem to be a trend or correlation between posterior predictive p-value and posterior probability of the clade *Photinus*. Thus, none of the chosen summary statistics is a good indicator of gene tree error (assuming the posterior probability of *Photinus* being monophyletic as a proxy for gene tree accuracy).

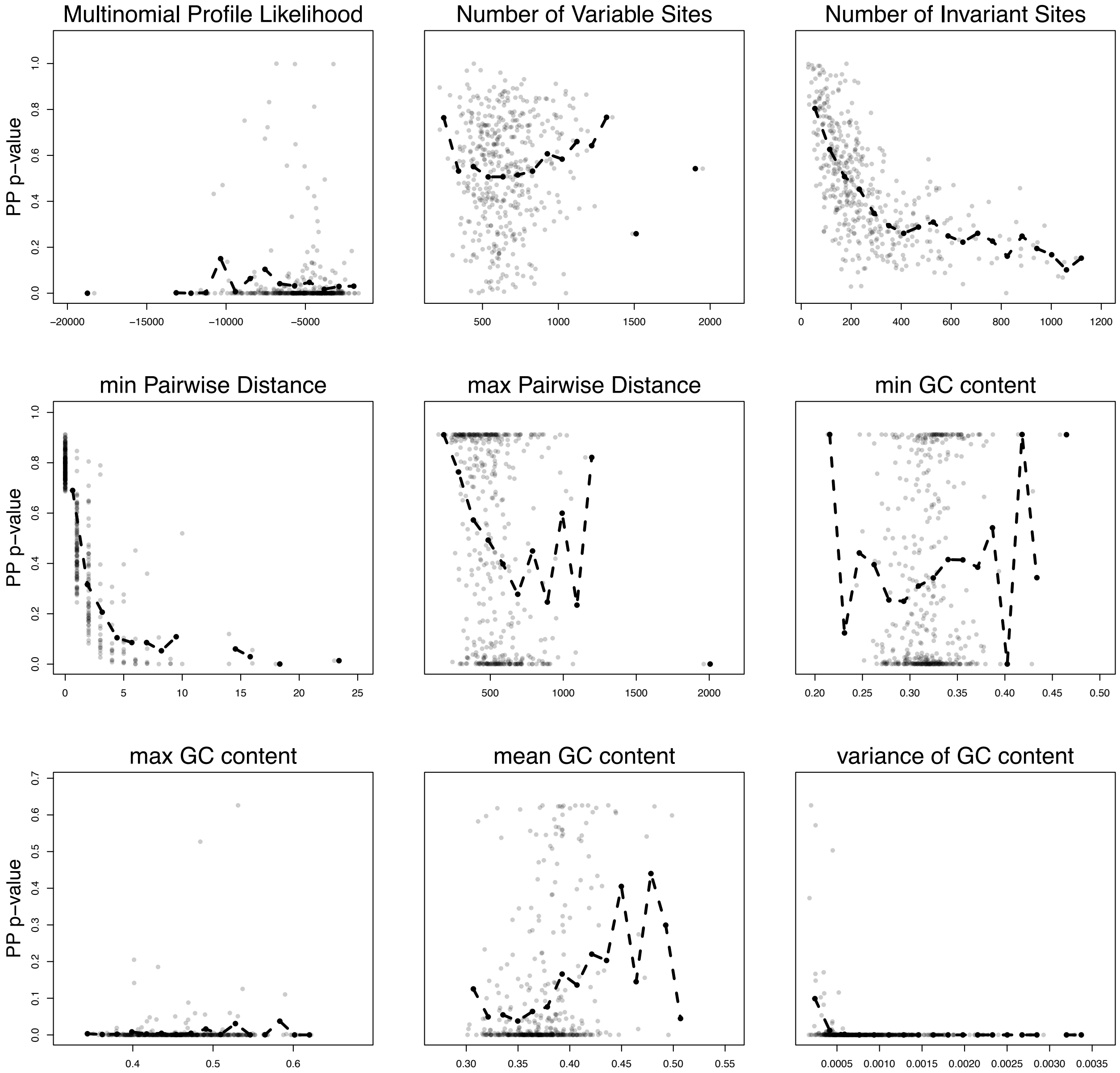


**Figure S11: Correlation between summary statistics and posterior predictive p-values.** The dashed black line shows a rolling average and the solid green line shows the regression line. There is a negative correlation between all summary statistics and the posterior predictive p-values. Model adequacy is thus improved when considering loci with lower values for these nine summary statistics.

### Gene tree discordance

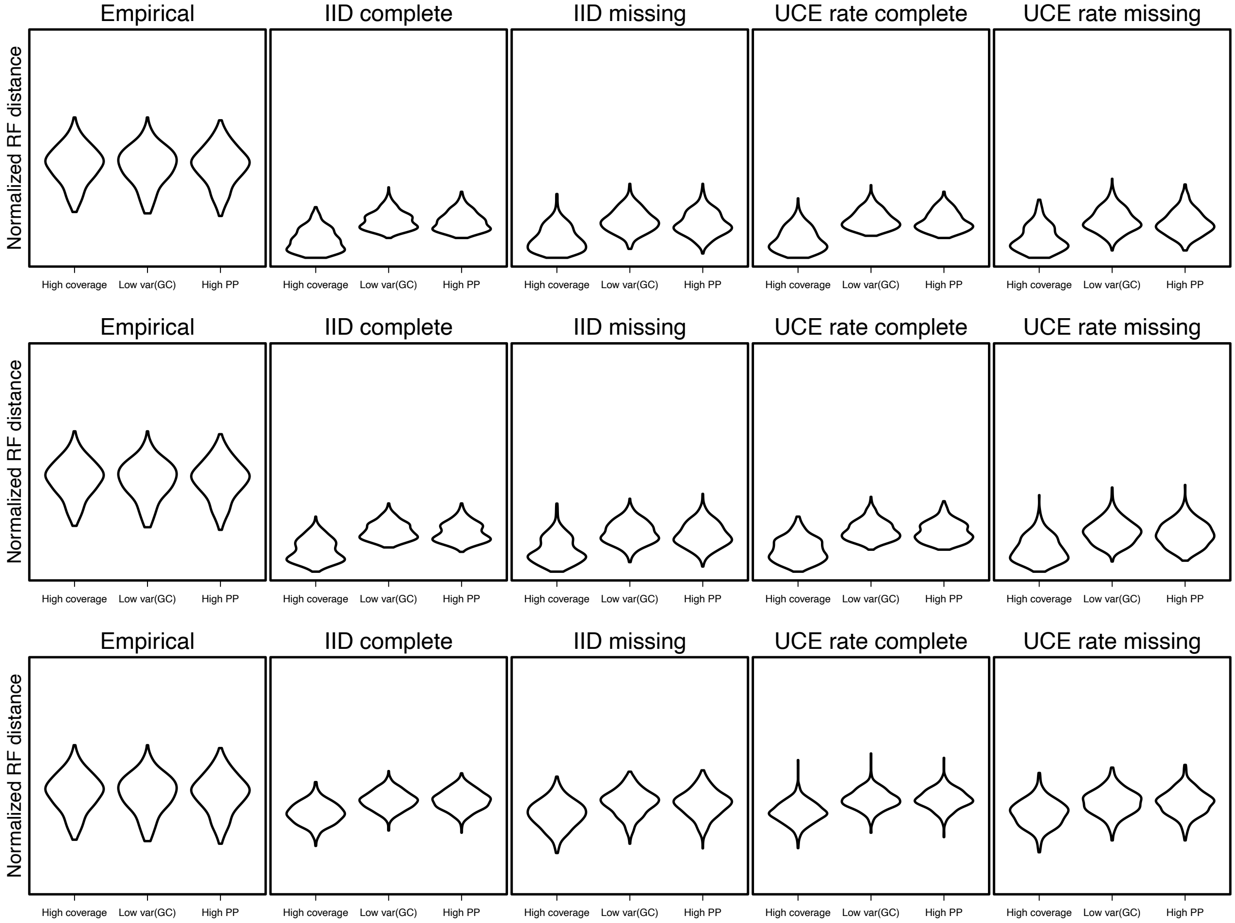


**Figure S12: Gene tree discordance measured using the normalized Robinson-Foulds (RF) distance between several reference trees and the single gene trees for all three simulation settings.** The top row shows the simulations for low populatio sizes (100,000), the middle row for intermediate population sizes (1,000,000) and the bottom row for large population sizes (10,000,000). As reference trees, we used the the maximum a posterior (MAP) phylogeny using the three different data subsets (high coverage, low variance in GC content, and high posterior probability of *Photinus+Ellychnia* being monophyletic). The left panel shows the frequency of the RF-distance for the empirical dataset of Martin et al. (2019). The second panel shows the RF-distance for the simulated dataset with complete sequences and homogeneous (*i.e.,* independent and identically distributed, IID) highly variables sites. The third panel shows the RF-distance for the simulated dataset with missing sequences and homogeneous highly variable sites. The fourth panel shows the RF-distance for the simulated dataset with complete sequences and systematically distributed (*i.e.,* akin to the empirical AHE dataset) highly variables sites. The right panel shows the RF-distance for the simulated dataset with missing sequences and systematically distributed highly variable sites. Neither of our simulation conditions are a strongly negative impact on gene tree discordance.

#### Posterior probabilities of given clades using single gene trees and the simulated data

**Table S2: Differences between Martin et al. (2019) phylogeny and the phylogeny estimated in this study.**

| Clade | Martin et al. (2019) | This study |
|---|---|---|
| Within the Lampyrinae | | |
| Ellychnia + Photinus pyralis | Monophyletic with 95% bootstrap support, but with a short branch between their shared ancestor to the next most recent common ancestor. | Ellychnia is not sister to Photinus pyralis, but diverges from Photinus macdermotti + Photinus brimleyi clade. |
| Photinus macdermotti 1 and 2 | | Flipped relative to Martin et. al |
| Ethra axillaris | Basal to Photinus + Pyractonema + Pyropyga | Basal to Photinus + Pyractonema + Pyropyga + Lucidota |
| Lucio blattinum | Sister to Lamprocera only, but with a short branch | Sister to Diaphnes + Lamprocera |
| Within the Photurinae | | |
| Photuris | Basal in the Photurinae | Derived within the Photurinae |
| Psilocladus | Sister to Photurinae + Amydetinae | Derived Within the Luciolinae |
| Within the Luciolinae | | |
| Luciola 3 + Emeia | Monophyletic, but with a short branch to the most recent common ancestor with the Pygoluciola + Luciola 4 + Curtos clade | Not monophyletic. Not sister to the Pygoluciola + Luciola 4 + Curtos clade |
| Asymmetricata + Luciola sp 2 | Basal to Emeia + Luciola + Pygoluciola + Curtos clade | |
| Pollaclassis | Sister to Pterotus, but with a short branch and relatively lower support | Basal to Pterotus |

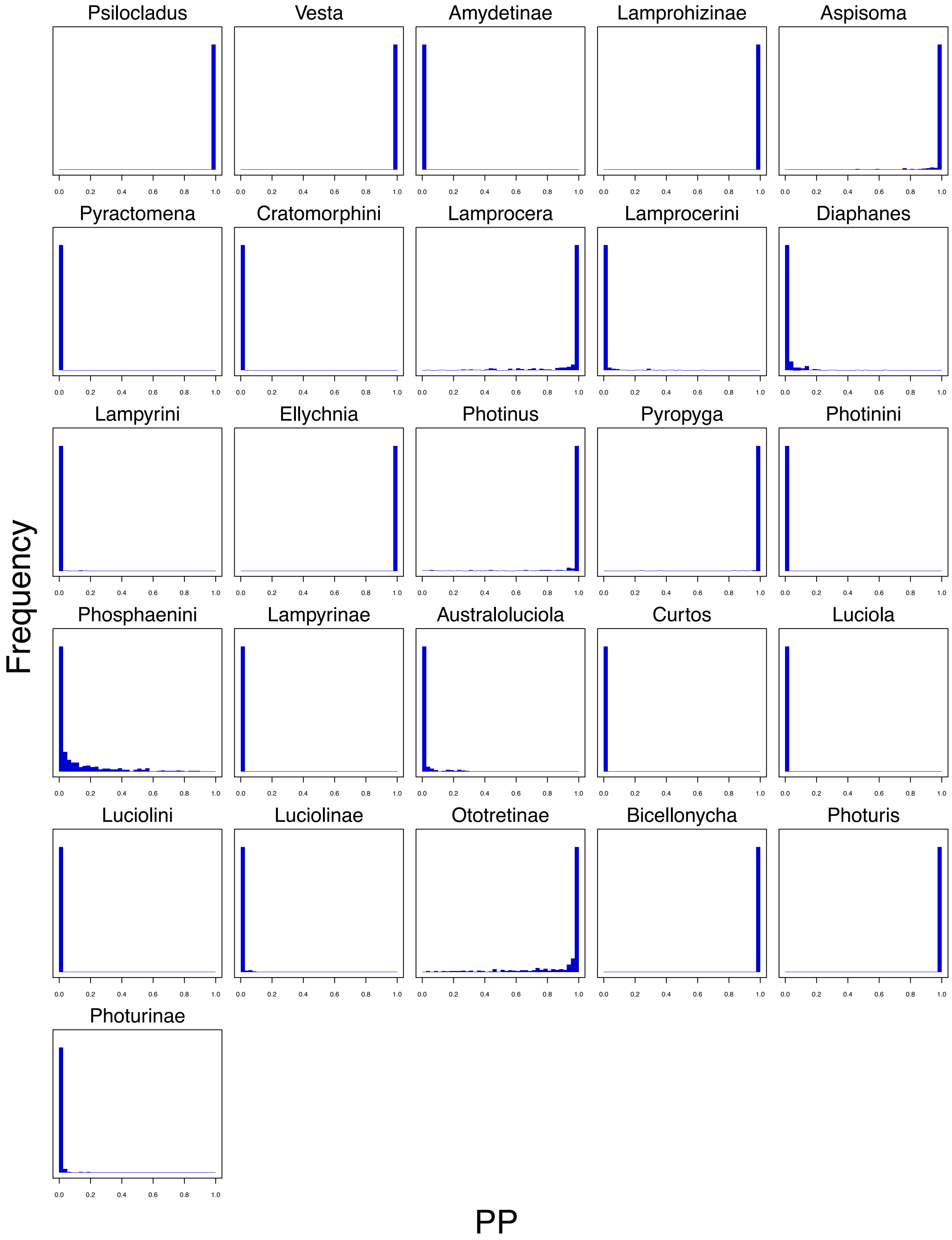


**Figure S13: Posterior probabilities for the clades using the simulated dataset with complete sequences and homogeneous (*i.e.,* independent and identically distributed, IID) highly variables sites.** The posterior probabilities without missing data, *i.e.,* complete sequences, show strong support for the correct clades.

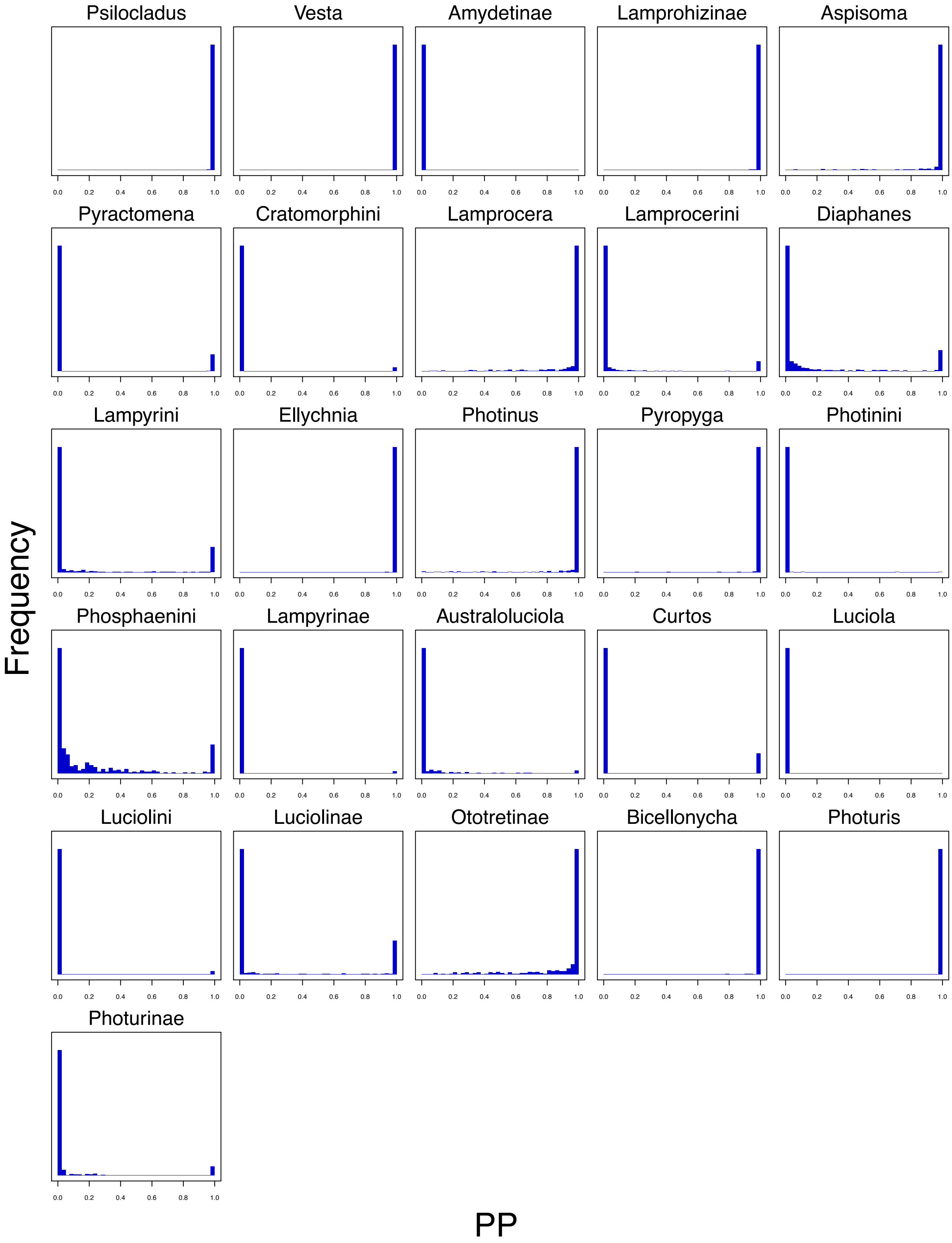


**Figure S14: Posterior probabilities for the clades using the simulated dataset with missing sequences and homogeneous (*i.e.,* independent and identically distributed, IID) highly variables sites.** The posterior probabilities with missing data has reduced support for the correct clade. Specifically, the *Photinus* clade has significantly reduced support, whereas Phosphaenini and *Australoluciola* have spuriously inflated support.

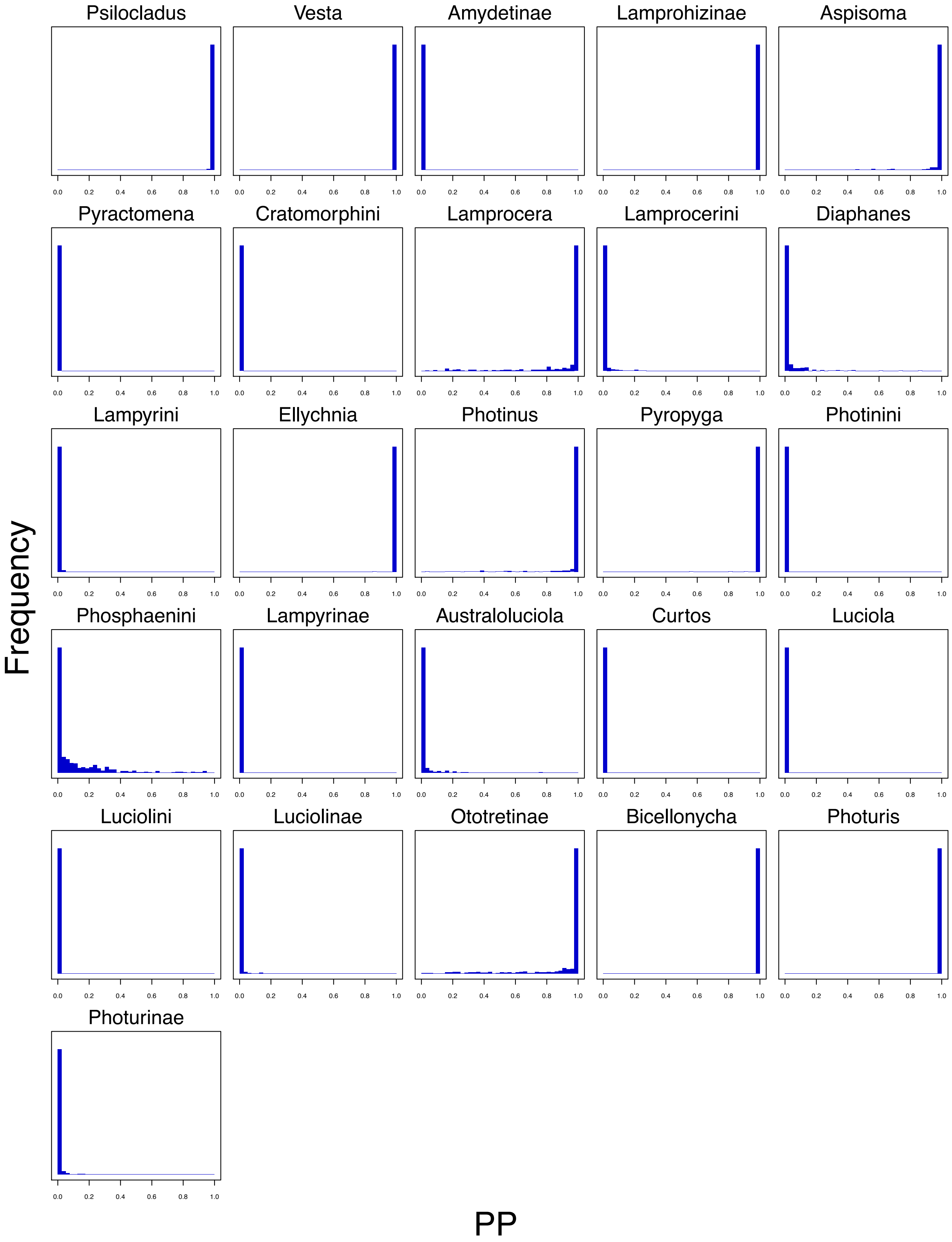


**Figure S15: Posterior probabilities for the clades using the simulated dataset with complete sequences and systematically distributed highly variables sites.** The posterior probabilities without missing data, *i.e.,* complete sequences, show strong support for the correct clades.

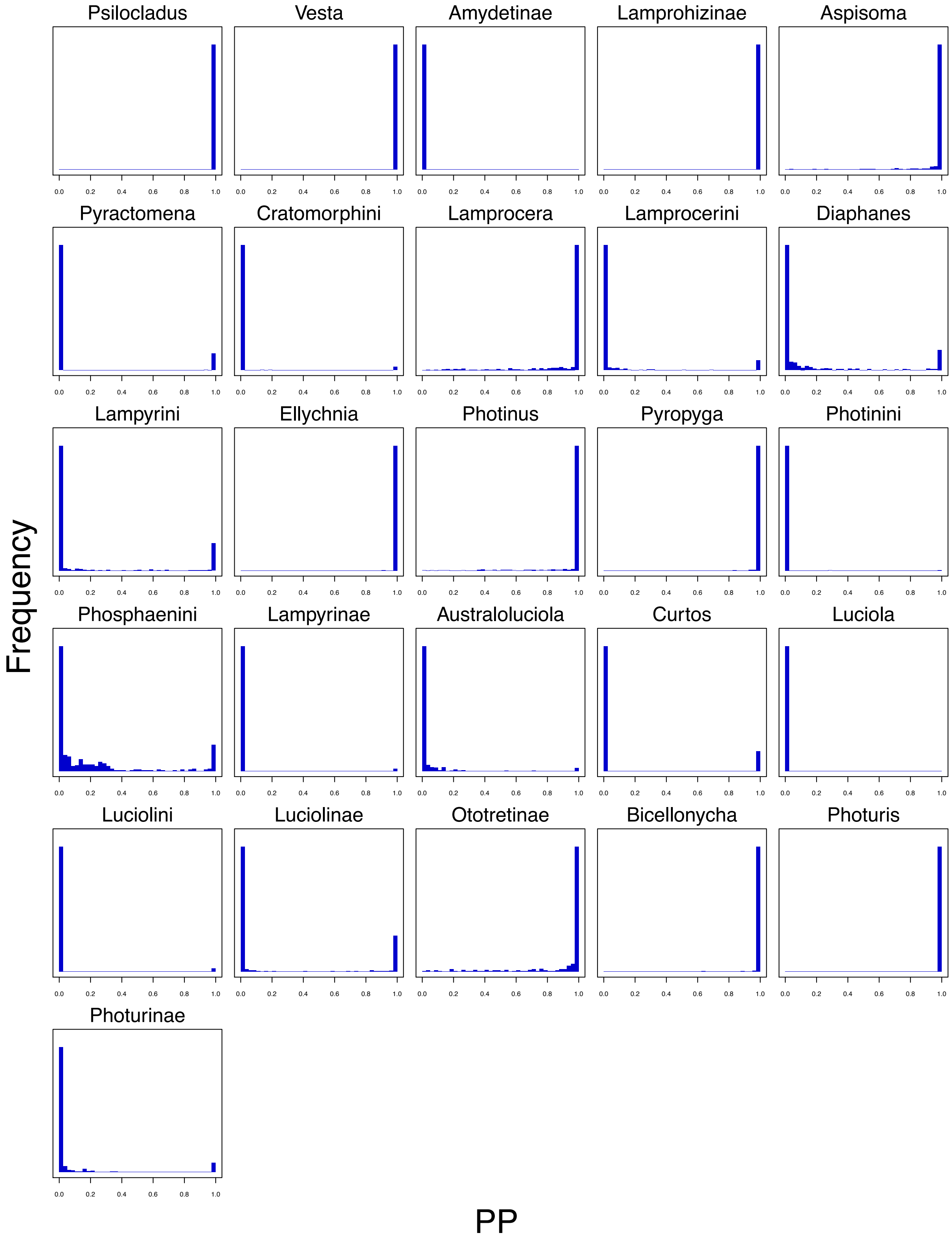


**Figure S16: Posterior probabilities for the clades using the simulated dataset with missing sequences and systematically distributed highly variables sites.** The posterior probabilities with missing data has reduced support for the correct clade. Specifically, the *Photinus* clade has significantly reduced support, whereas Phosphaenini and *Australoluciola* have spuriously inflated support.

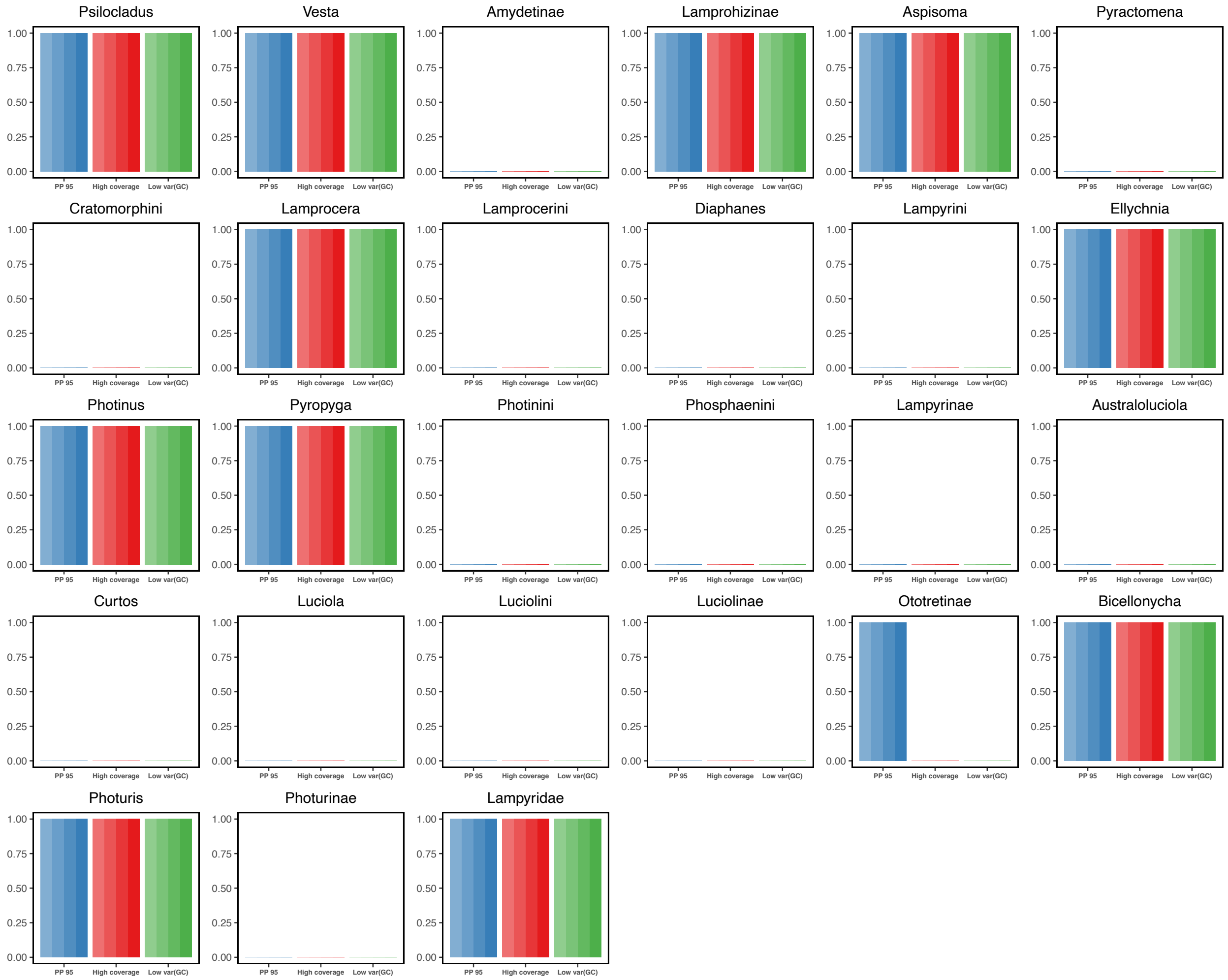


**Figure S17: Posterior probabilities of the different clades being monophyletic for the three data subsets.** The results show the posterior probabilities of the four MCMC replicates (different color shades) for the three data subsets: High posterior probability of *Photinus* being monophyletic (blue), high sequence coverage (red) and low variance in GC content (green). The analyses produced identical posterior probabilities for all clades except Ototretinae. These posterior probabilities for all clades show extreme certainty; either full support with a posterior probability of 1.0 or full rejection with a posterior probability of 0.0.

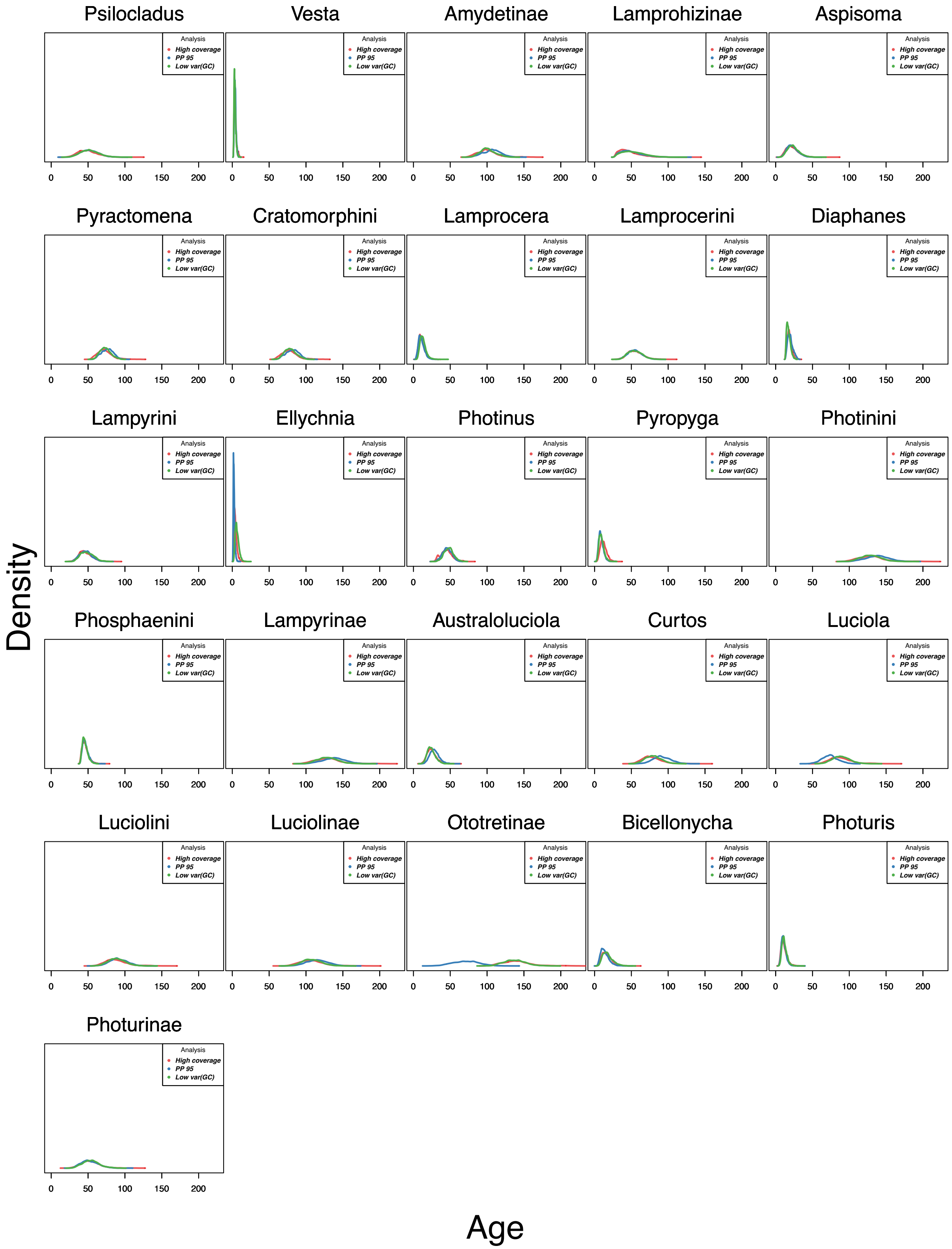


**Figure S18: Crown age estimates of the named clades from Table S1.** Each plot shows the posterior distribution of the crown age of a different clade for the three data subsets: High posterior probability of *Photinus* being monophyletic (blue), high sequence coverage (red) and low variance in GC content (green). The estimated ages agree between the three data subsets except for the cases when the topology differed.
